# Targeting a Therapy-Resistant Cancer Cell State Using Masked Electrophiles as GPX4 Inhibitors

**DOI:** 10.1101/376764

**Authors:** John K. Eaton, Laura Furst, Richard A. Ruberto, Dieter Moosmayer, Roman C. Hillig, André Hilpmann, Katja Zimmermann, Matthew J. Ryan, Michael Niehues, Volker Badock, Anneke Kramm, Sixun Chen, Paul A. Clemons, Stefan Gradl, Claire Montagnon, Kiel E. Lazarski, Sven Christian, Besnik Bajrami, Roland Neuhaus, Ashley L. Eheim, Vasanthi S. Viswanathan, Stuart L. Schreiber

## Abstract

We recently discovered that inhibition of the lipid peroxidase GPX4 can selectively kill cancer cells in a therapy-resistant state through induction of ferroptosis. Although GPX4 lacks a conventional druggable pocket, covalent small-molecule inhibitors are able to overcome this challenge by reacting with the GPX4 catalytic selenocysteine residue to eliminate enzymatic activity. Unfortunately, all currently-reported GPX4 inhibitors achieve their activity through reactive chloroacetamide groups. We demonstrate that such chloroacetamide-containing compounds are poor starting points for further advancement given their promiscuity, instability, and low bioavailability. Development of improved GPX4 inhibitors, including those with therapeutic potential, requires the identification of new electrophilic chemotypes and mechanisms of action that do not suffer these shortcomings. Here, we report our discovery that nitrile oxide electrophiles, and a set of remarkable chemical transformations that generates them in cells from masked precursors, provide an effective strategy for selective targeting of GPX4. Our results, which include structural insights, target engagement assays, and diverse GPX4-inhibitor tool compounds, provide critical insights that may galvanize development of improved compounds that illuminate the basic biology of GPX4 and therapeutic potential of ferroptosis induction. In addition, our discovery that nitrile oxide electrophiles engage in highly selective cellular interactions and are bioavailable in their masked forms may be relevant for targeting other currently undruggable proteins, such as those revealed by recent proteome-wide ligandability studies.

## INTRODUCTION

We have previously reported that cancer cells in a drug-induced, therapy-resistant state have an enhanced dependence on the lipid peroxidase activity of GPX4 to prevent undergoing ferroptotic cell death^1,2^. *In vitro* and *in vivo* evidence support the possibility that rational combination of GPX4 inhibitors with standard-of-care regimens based on chemotherapy, targeted therapy^1,2^ and immunotherapy^3^ can achieve durable responses in a range of cancer types.

GPX4 presents a challenge for the development of effective small-molecule inhibitors due to the relatively flat surface adjacent to its key catalytic selenocysteine residue. Indeed, the only cell-active small molecules that have been shown to inhibit GPX4 directly act by covalently binding the selenocysteine residue via a reactive α-chloroacetamide group^4,5^. Major limitations of such chloroacetamide GPX4 inhibitors include their off-target effects and poor stability, detracting from their use as tools to study GPX4 biology *in vitro* and explore therapeutic hypotheses *in vivo*. We have therefore sought to identify new structural classes of GPX4 inhibitors that address the limitations of the existing chloroacetamide class and can serve as starting points for the development of improved compounds.

## RESULTS

### ML210 exhibits cellular activity similar to known chloroacetamide GPX4 inhibitors

To identify small-molecule GPX4 inhibitors that do not contain a chloroacetamide warhead, we focused our attention on the nitroisoxazole-containing compound ML210 (Figure 1A)^6^. Although ML210 (Figure 1A) lacks an obvious protein-reactive group through which it could act as a GPX4 inhibitor, data from the Cancer Therapeutics Response Portal (CTRP at http://portals.broadinstitute.org/ctrp.v2.1/)^7–9^ reveal that it exhibits a pattern of cell killing across 821 cancer cell lines that is strikingly similar to that of chloroacetamide GPX4 inhibitors (1*S*,3*R*)-RSL3 (RSL3) and ML162 (Figures 1A and 1B). We further confirmed previous reports that GPX4 enzymatic activity is eliminated in cells treated with ML210^1,4^ (Figure 1C), an observation that is consistent with, though not conclusive of, ML210 acting as a GPX4 inhibitor. Like known GPX4 inhibitors, ML210 treatment of cells results in the accumulation of cellular lipid hydroperoxides whose lethal effects can be prevented by co-treatment with ferrostatin-1 (fer-1) (Figure 1D). Fer-1 is a lipophilic radical-trapping antioxidant that specifically rescues cellular effects caused by the loss of GPX4 activity^10–12^. Notably, fer-1 rescues completely the cell-killing activity of ML210 but not that of either RSL3 or ML162 (Figure 1E), suggesting that if ML210 is indeed a direct inhibitor of GPX4, it could prove much more selective than chloroacetamide inhibitors (Figure 1E). Together, these observations encouraged us to further explore ML210 as a potential direct and selective inhibitor of GPX4 with a novel mechanism of action.

**Figure 1.**
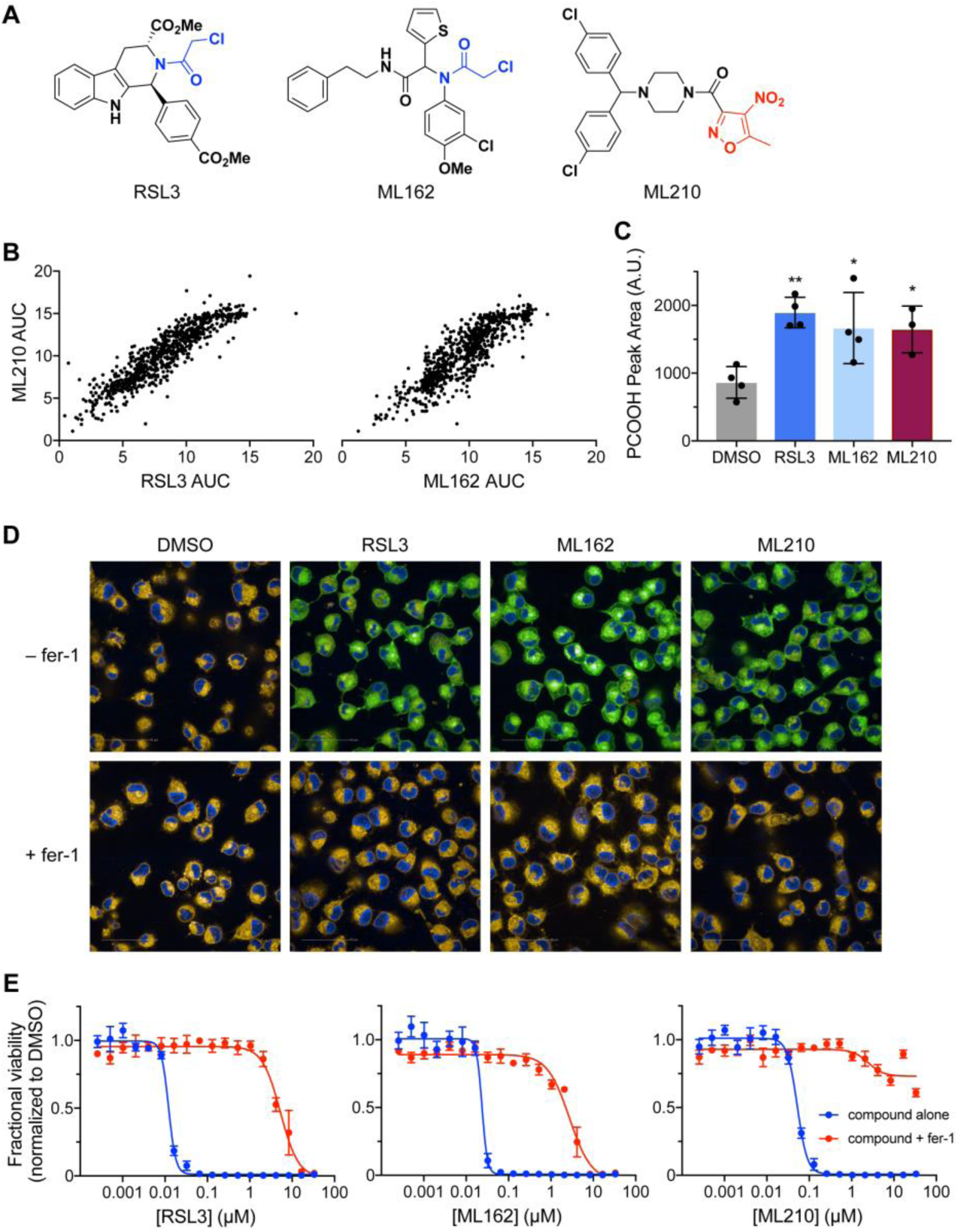
ML210 exhibits cell-killing activity similar to known GPX4 inhibitors. (A) Chemical structures of RSL3, ML162, and ML210. Chloroacetamide groups shown in blue, nitroisoxazole group shown in red. (B) ML210 exhibits cell-killing activity similar to RSL3 and ML162 across a panel of 821 cancer cell lines. Each dot represents a single cancer cell line. Data plotted reflect an area-under-the-curve (AUC) metric of cell-line sensitivity to small molecules. (C) Treatment of LOX-IMVI cells with RSL3, ML162, or ML210 inhibits the reduction of GPX4-specific substrate phosphatidylcholine hydroperoxide (PCOOH) in cell lysates. Data are plotted as the mean ± s.d., n ≥ 3 technical replicates. \**p* < 0.05, *\*\*p <* 0.005 vs. DMSO control. Scale bars: 50 μm. (D) Treatment of cells with RSL3, ML162, or ML210 leads to accumulation of lipid peroxides as assessed by C11-BODIPY 581/591 fluorescence in LOX-IMVI cells. The C11-BODIPY dye emission shifts from orange to green upon oxidation. Co-treatment with lipophilic antioxidant ferrostatin-1 (fer-1) prevents compound-induced lipid peroxide accumulation. (E) Co-treatment with fer-1 rescues the cell-killing effects of RSL3, ML162, and ML210 in LOX-IMVI melanoma cells. Data are plotted as mean ± s.e.m., n = 4 technical replicates.

### ML210 is a selective covalent inhibitor of cellular GPX4

To test whether ML210 interacts directly with cellular GPX4 in a manner similar to chloroacetamide GPX4 inhibitors, we developed alkyne-containing analogs of ML210, RSL3, and ML162. These alkyne probes exhibit cellular activity similar to their respective parent compounds (Figures 2A and S1A) and enable affinity enrichment experiments that can be used to identify the covalent protein targets of these compounds (see Extended Methods). We validated this system using chloroacetamide probes RSL3-yne and ML162-yne, both of which were observed to pull down GPX4 from cells (Figure 2B). The ML210 analog ML210-yne similarly pulls down GPX4 from cells, confirming ML210 as a direct and covalent inhibitor of GPX4 (Figure 2B).

**Figure 2.**
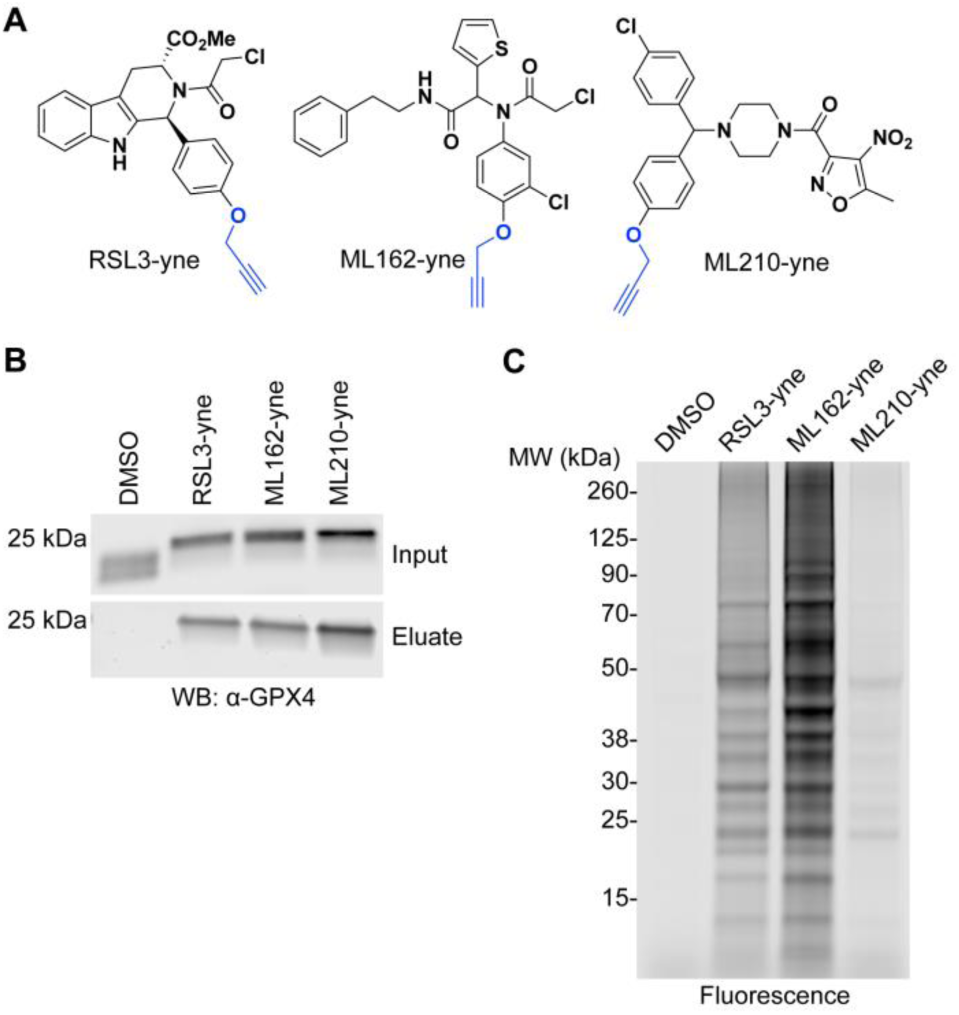
ML210 is a selective, covalent inhibitor of cellular GPX4. (A) Structures of GPX4-inhibitor affinity probes RSL3-yne, ML162-yne, and ML210-yne with alkyne groups shown in blue. (B) RSL3-yne, ML162-yne, and ML210-yne pull down GPX4 from cells. (C) Fluorescent labeling of proteins modified by RSL3-yne, ML162-yne, and ML210-yne probes in LOX-IMVI cells.

These same alkyne probes can also be used to assess the proteome-wide selectivity of covalent GPX4 inhibitors. A major drawback of chloroacetamide GPX4 inhibitors is their high chemical reactivity, which imparts them with low proteome-wide selectivity. To test whether ML210 acts in a more selective manner, we performed proteome reactivity profiling experiments using the alkyne probes described above. ML210-yne exhibits markedly lower proteome reactivity compared to the RSL3 and ML162 probes, which engage in a large number of covalent interactions across the proteome (Figure 2C and S1B). Competitive labeling experiments indicate that the off-target proteins of ML210 are shared with chloroacetamide GPX4 inhibitors (Figures S1C and S1D). Mass spectrometry-based proteomics reveals that these off-targets are highly abundant proteins (e.g. tubulins) (Figure S1E), which are targeted by ML210 only at high concentrations and impact only a minor fraction of each protein’s pool.

Together, these results establish ML210 as a potent and selective direct inhibitor of GPX4 with compelling properties that overcome the critical shortcomings of chloroacetamide inhibitors. Specifically, despite having a covalent mechanism of action, which may be necessary for targeting GPX4, ML210 is highly selective as evidenced by low proteome-wide reactivity and full rescuability by fer-1 (Figure 1E). This selectivity appears to be related to the nitroisoxazole group of ML210. We found that replacement of the nitroisoxazole warhead with either chloroacetamide or propiolamide groups results in analogs that are less potent and significantly less selective (Figure S2A), as the effects of these compounds can be rescued only partially by fer-1. In contrast, the selectivity of chloroacetamide GPX4 inhibitors can be improved in certain cases by exchanging the chloroacetamide moiety for a nitroisoxazole warhead (Figures S2B and S2C).

### ML210 is a prodrug that requires cellular activation

To understand the ML210-GPX4 interaction, we developed an intact protein mass spectrometry workflow to detect covalent GPX4-inhibitor complexes generated in cells. We tested this system using chloroacetamide GPX4 inhibitors RSL3 and ML162, and found that they both form covalent adducts with GPX4 in cells (Figures 3A, S3A, and S3B). Subjecting ML210 to this approach also revealed the formation of a covalent adduct with GPX4 (Figure 3A). This confirms, once again, that ML210 is a covalent GPX4 inhibitor. However, the ML210-bound GPX4 adduct mass (434 Da) corresponds to an unusual 41 Da loss from ML210 that cannot be explained by the displacement of an obvious leaving group and suggests a novel covalent mechanism of action for ML210.

**Figure 3.**
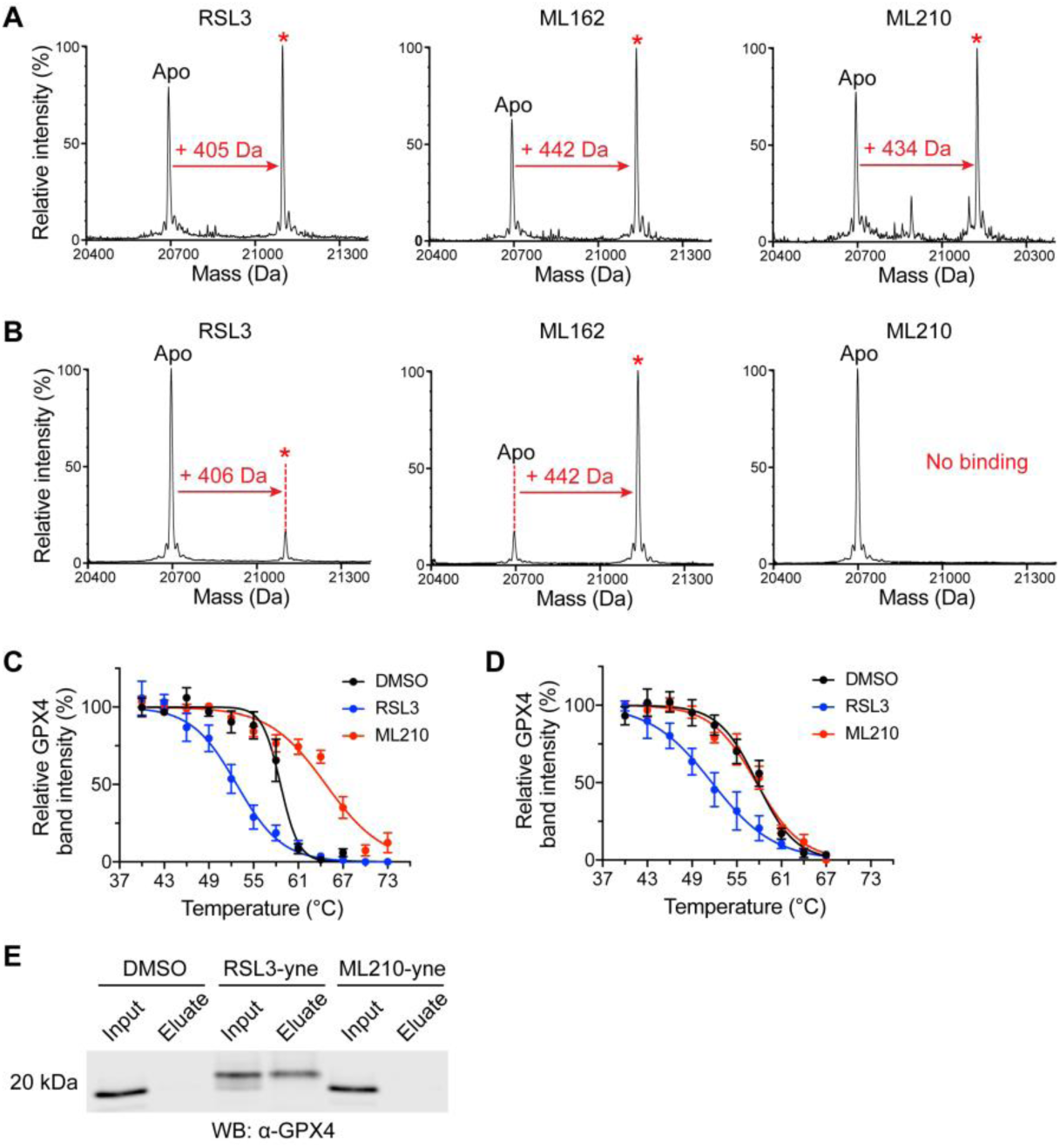
ML210 is a prodrug that requires cellular activation to bind GPX4. (A) RSL3, ML162, and ML210 show evidence of covalent interaction with cellular GPX4 by intact protein mass spectrometry. Covalent adduct peaks are marked with a red asterisk (*). See also Figure S3B. (B) RSL3 and ML162, but not ML210, covalently bind purified GPX4 as assessed by intact protein mass spectrometry. Adduct peaks are marked with a red asterisk (*). (C) GPX4 CETSA profiles for intact cells treated with DMSO (black), RSL3 (blue), or ML210 (red). Data are plotted as mean ± s.e.m., n ≥ 4 biologically independent samples. (D) GPX4 CETSA profiles for cell lysates treated with DMSO (black), RSL3 (blue), or ML210 (red). Data are plotted as mean ± s.e.m., n = 3 biologically independent samples. (E) RSL3-yne, but not ML210-yne, pulls down GPX4 from cell lysate.

We sought to explore this further by reacting ML210 biochemically with purified GPX4 protein. From this experiment, we discovered that the ML210-GPX4 adduct forms only when ML210 is added to intact cells and not when ML210 is incubated with purified wild type GPX4 (Figure 3B). This context-dependent binding is in contrast to chloroacetamide inhibitors, which form adducts with GPX4 both when added to cells or incubated with purified protein (Figure 3B).

To confirm the intriguing intact-cell dependence of ML210 binding, we developed two additional assays that report on cellular GPX4 target engagement. The first method, a label-free cellular thermal shift assay (CETSA), allows for protein thermal stability to be assessed in cells or cell lysates using western blot detection^13^. Consistent with mass spectrometry-based binding experiments, CETSA reveals that ML210 engages GPX4 in intact cells but not cell lysates, while chloroacetamide inhibitors interact with GPX4 in both contexts (Figures 3C, 3D, and S3C). CETSA also revealed another striking distinction between ML210 and chloroacetamide GPX4 inhibitors: ML210 substantially stabilizes GPX4 while chloroacetamide inhibitors result in GPX4 thermal destabilization. This contrast may reflect underlying differences in the way these two classes of molecules interact with GPX4.

The second GPX4 target-engagement method we used is a GPX4 pulldown assay that relies on the previously described alkyne-tagged analogs of ML210, RSL3, and ML162 (Figure 2A). Consistent with our other results, assessment of GPX4 inhibitors with this cellular target engagement assay confirmed that ML210, but not chloroacetamide inhibitors, requires intact cells to engage GPX4 covalently (Figure 3E).

The intact-cell dependence of ML210 together with its lack of an obvious protein-reactive moiety led us to hypothesize that ML210 is a prodrug that undergoes chemical transformation in cells to covalently modify GPX4 via a novel electrophile. Such a mechanism could also explain the unusual mass of the ML210-GPX4 adduct, which may be a result of mass loss during cellular activation.

### ML210 binds the GPX4 selenocysteine residue

As a first step toward understanding how a cellular metabolite of ML210 may bind and inhibit GPX4, we performed experiments to identify the residue of GPX4 that is covalently modified by ML210. Chloroacetamide inhibitors have been reported to bind GPX4 only when the active site contains a selenocysteine or cysteine residue at position 46^5,14^. By obtaining, for the first time, a co-crystal structure of GPX4 in complex with ML162 (Figure 4A; Table S1; see Extended Discussion), we now demonstrate conclusively that the GPX4 selenocysteine residue is indeed the target of chloroacetamide GPX4 inhibitors. We were unable to use this same approach to characterize ML210 binding because of the inability of ML210 per se to interact with purified GPX4 protein and difficulties associated with scaling in-cell binding preparations. Instead, to identify the GPX4 residue that ML210 covalently modifies, we performed competitive pulldown experiments with ML210-yne and chloroacetamide inhibitors (Figure 4B). These experiments revealed that the binding of ML210 and chloroacetamide inhibitors is mutually exclusive, signifying that ML210 likely binds the GPX4 selenocysteine residue.

**Figure 4.**
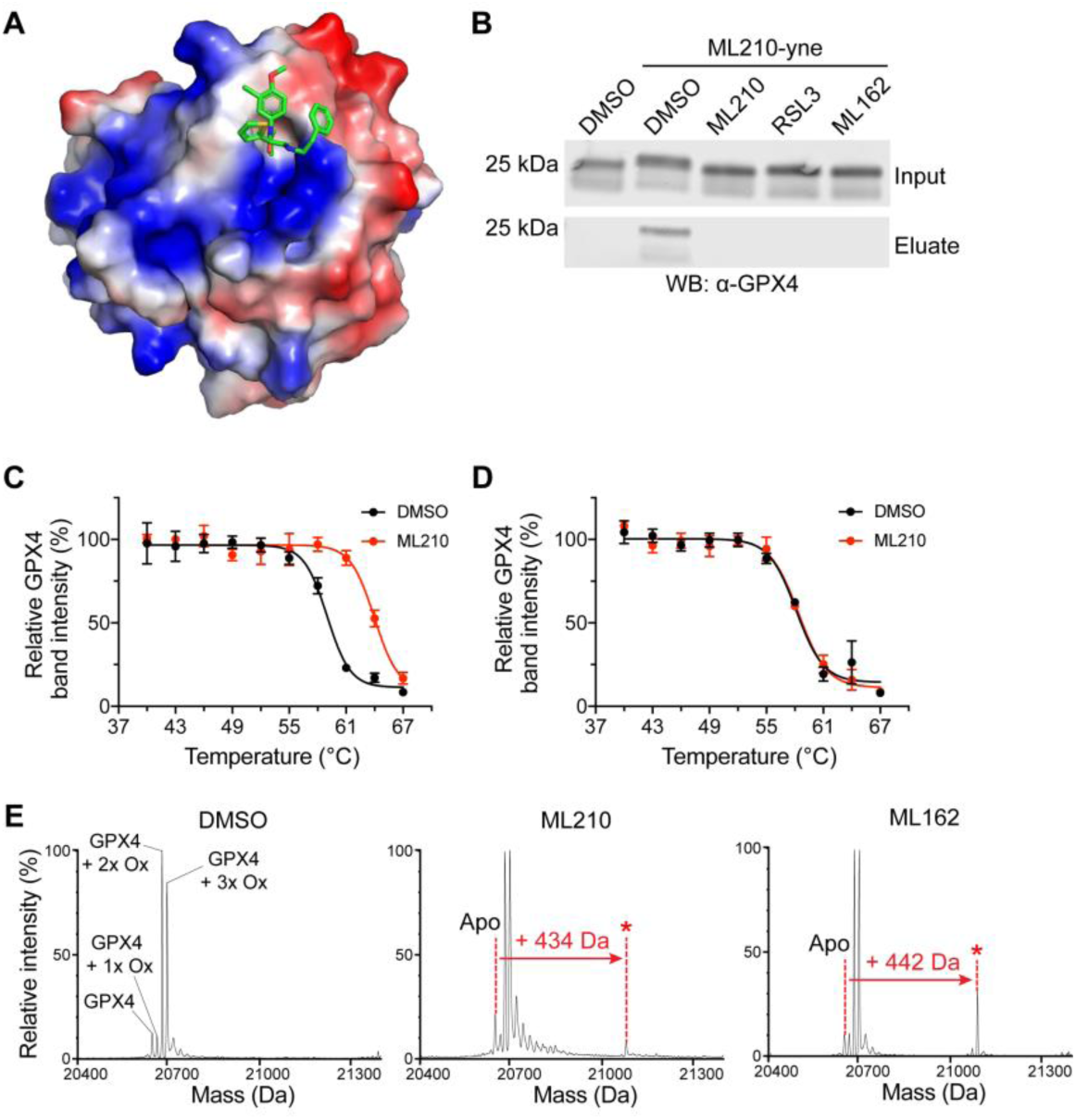
ML210 and chloroacetamide inhibitors covalently bind GPX4 selenocysteine residue. (A) Co-crystal structure of GPX4^C66S^ bound to *S*-enantiomer of ML162. (B)Competitive affinity enrichment between ML210-yne probe and RSL3 and ML162. (C) G PX4 CETSA of intact cells expressing GPX4^U46C^-mutant protein treated with DMSO (black) or ML210 (red). Data are plotted as mean ± s.e.m., n = 3 biologically independent samples. (D) GPX4 CETSA of intact cells expressing GPX4^U46A^-mutant protein treated with DMSO (black) or ML210 (red). Data are plotted as mean ± s.e.m., n = 3 biologically independent samples. (E) ML210 and ML162 interact with GPX4^U46C^ protein in cells. Compared to wild type GPX4, GPX4^U46C^ is observed in multiple oxidation states. See Figure S3D for RSL3 binding data.

To further confirm the role of the selenocysteine in ML210 binding, we performed intact-cell CETSA experiments with GPX4 mutants containing alanine or cysteine in place of the catalytic selenocysteine residue. We found that both chloroacetamide inhibitors and ML210 bind the cysteine-mutant form of GPX4 (Figures 4C and 4D). However, neither inhibitor class interacts with the alanine-mutant protein (Figure 4D). We confirmed inhibitor binding to GPX4^U46C^ by intact protein mass spectrometry (Figure 4E and S3D). These findings establish the importance of a nucleophilic (seleno)cysteine residue within the active site of GPX4 for covalent interaction with inhibitors. Moreover, these data indicate that GPX4 inhibitors do not covalently bind other nucleophilic residues of GPX4 in cells.

Together, these observations demonstrate that the greater selectivity of ML210 compared to chloroacetamide inhibitors is not due to covalent binding with a different GPX4 residue. Rather, the basis of ML210’s selectivity may be a combination of its prodrug nature and the unique reactivity of its unleashed, active electrophilic form.

### Cellular activation of ML210 generates the α-nitroketoxime JKE-1674

We next aimed to identify the ML210-derived electrophile formed in cells that targets GPX4. The nitroisoxazole group contains structural features that are susceptible to diverse chemical transformations in cells^15–17^. To understand the role that this functional group may play in the mechanism of action of ML210, we performed structure-activity relationship (SAR) studies (Figures 5A and S4A). These revealed that both the nitro and isoxazole groups are unable to be substituted without complete loss of activity (Figure 5A)^6^. In contrast to this steep SAR of the nitroisoxazole, modifications of substituents at the 3-and 5-positions of the isoxazole ring are well tolerated with little effect on potency relative to ML210 (Figures 5A and S4A). Most surprisingly, SAR of the nitroisoxazole 5-position revealed that cellular activity is not affected by steric bulk or polarity of substituents (Figure S4). Together with the unusual mass loss associated with ML210 binding to GPX4 (Figure 3A), this SAR led us to hypothesize that cellular activation of ML210 may involve elimination of the nitroisoxazole 5-position. Consistent with this hypothesis, a 5-isopropyl ML210 analog (Figure 5B) gives rise to a cellular GPX4 adduct of identical mass to the ML210-derived GPX4 adduct (Figure 5C).

**Figure 5.**
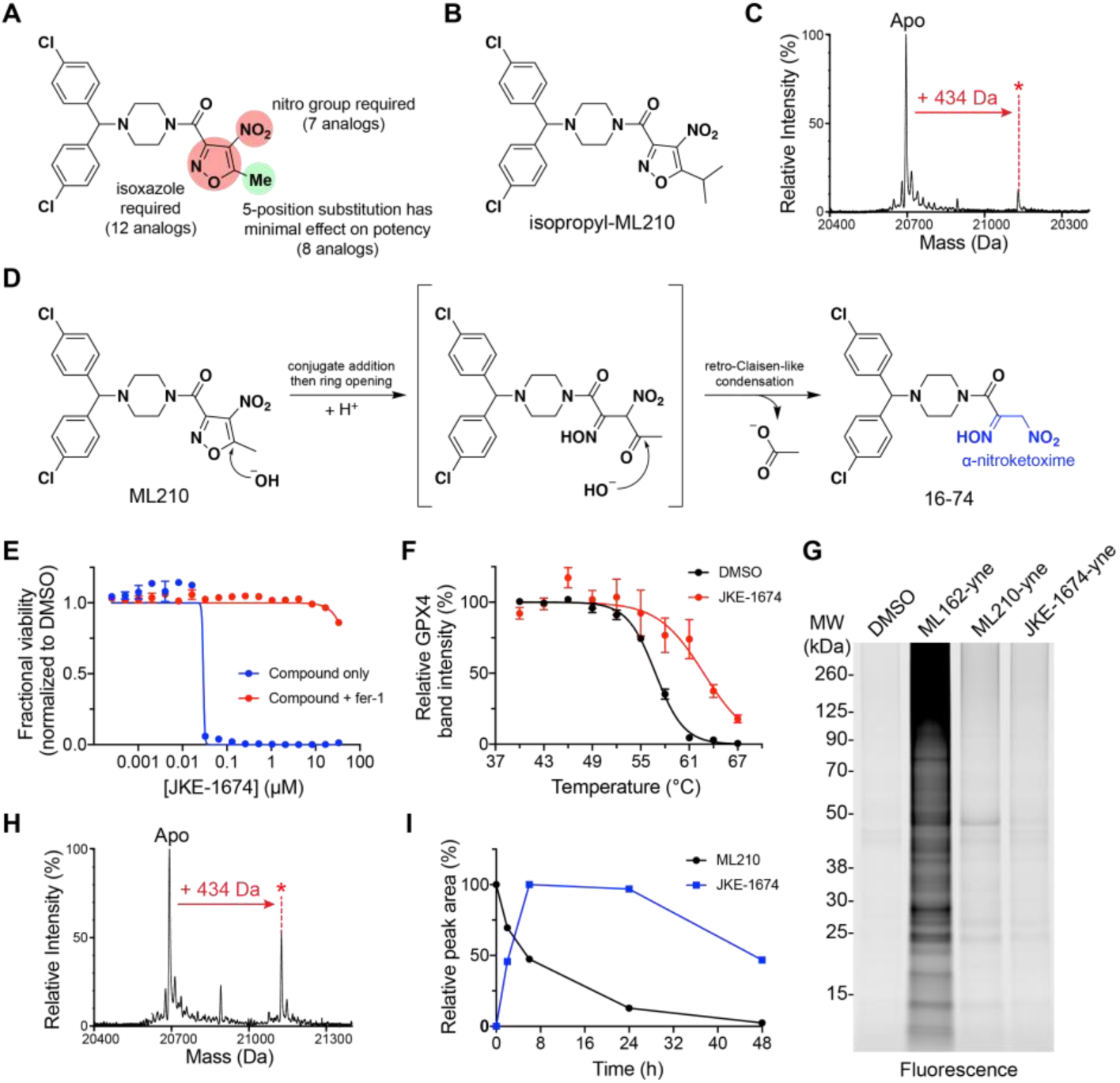
Cellular activation of ML210 generates α-nitroketoxime JKE-1674. (A) Summary of ML210 nitroisoxazole SAR studies. See also Figure S4. (B) Structure of isopropyl-ML210. (C) Isopropyl-ML210 produces the same covalent adduct mass (434 Da) with cellular GPX4 as ML210. (D) Scheme showing proposed ML210 hydrolysis and structure of JKE-1674 with α-nitroketoxime group highlighted in blue. (E) Co-treatment with fer-1 rescues the LOX-IMVI cell-killing effects of JKE-1674 and ML210 to a similar extent. Data are plotted as mean ± s.e.m., n = 4 technical replicates. (F) GPX4 CETSA of intact cells treated with JKE-1674 (red) reveals thermal stabilization of GPX4 compared to treatment DMSO (black). Data are plotted as mean ± s.e.m., n ≥ 4 biologically independent samples. (G) A JKE-1674 alkyne analog (JKE-1674-yne, see Figure S5A) exhibits proteome-wide reactivity similar to ML210-yne. (H) JKE-1674 produces a GPX4 covalent adduct of the same mass as the ML210-GPX4 covalent adduct (434 Da). (I) JKE-1674 forms in a time-dependent manner in cells treated with ML210. See also Figure S5.

Nitroisoxazoles have been reported to undergo base-promoted hydrolysis to yield an α-nitroketoxime and a carboxylic acid^18^. By investigating the relevance of this reaction to ML210 activation, we identified aqueous basic conditions that rapidly convert ML210 to its corresponding α-nitroketoxime (Figure 5D).

This α-nitroketoxime, JKE-1674 (Figure 5D), exhibits GPX4 inhibitory activity that is indistinguishable from that of ML210 in LOX-IMVI viability and CETSA experiments (Figures 5E and 5F). An alkyne analog of JKE-1674 (Figure S5A) also pulls down GPX4 from cells (Figure S5C) and retains the selective proteome-wide reactivity profile of ML210 (Figure 5G). Notably, JKE-1674 yields the same 434 Da covalent GPX4 adduct in cells as ML210 and isopropyl-ML210 (Figure 5H). We further confirmed a mechanistic link between JKE-1674 and ML210 by observing the time-dependent formation of JKE-1674 in cells treated with ML210 (Figures 5I, S5D, and S5E).

Together, these data provide strong evidence that JKE-1674 is a key metabolite in the mechanism of action of ML210. However, like ML210, JKE-1674 is not able to engage either purified recombinant GPX4 or GPX4 in cell lysate (Figures S5I and S5J), indicating that additional cellular activation of JKE-1674 is required to yield a protein-reactive electrophile.

### Dehydration of JKE-1674 yields a nitrile oxide electrophile that binds GPX4

The mass of the ML210-derived GPX4 adduct suggests that the final protein-reactive ML210 metabolite is related to JKE-1674 through the loss of water. Such dehydration of JKE-1674 could lead to several species capable of acting as electrophiles (Figure S6A). To synthesize and interrogate the relevance of these different metabolites for GPX4 binding, we attempted to identify reaction conditions that eliminate water from JKE-1674. Unfortunately, we found that published procedures for the dehydration of α-nitroketoximes do not promote this transformation with JKE-1674^19,20^. However, we developed a two-step reaction sequence (Figure S6B) that enables the isolation of a dehydrated analog of JKE-1674, JKE-1777 (Figure 6A), which exhibits an infrared absorbance at 2286 cm^-1^ characteristic of a nitrile oxide group^21,22^.

JKE-1777 is unstable in solution, precluding detailed characterization, but it forms dimeric degradation products consistent with nitrile oxide reactivity (Figure S6C)^21^. We found that JKE-1777 exhibits cell killing that is rescuable by fer-1 (Figure 6B). The instability of JKE-1777 may explain its diminished potency compared to other ML210-derived compounds, as the degradation products of JKE-1777 are inactive (Figures S6C and S6D). Critically, whereas ML210 and JKE-1674 cannot react covalently with purified GPX4, JKE-1777 can react with purified GPX4 and yields the same 434 Da adduct that other ML210-derived analogs generate with GPX4 only in cells (Figure 6C). Based on these observations, we hypothesized that the nitrile oxide JKE-1777 is the ultimate ML210-derived electrophile that targets GPX4 in cells.

To corroborate the role of a nitrile oxide electrophile in inhibiting GPX4, we attempted to synthesize JKE-1777 analogs containing stable nitrile oxide precursors. We were unable to prepare an analog containing a hydroximoyl chloride group, which is commonly used in chemical synthesis for the *in situ* preparation of nitrile oxides^23^. Instead, we found that JKE-1674 can be converted directly into JKE-1708 (Figures 6D and S7A), which contains the related nitrolic acid group that can also act as a nitrile oxide precursor^24^. Functional assessment of JKE-1708 revealed cell-killing effects that are fully rescued by fer-1 in a manner similar to rescue of ML210 and JKE-1674 (Figures 6E and S7C). Analysis of JKE-1708 by CETSA revealed thermal stabilization of GPX4 identical to that induced by ML210 and JKE-1674 (Figure 6F). Like JKE-1777, JKE-1708 similarly does not require cellular activation and reacts spontaneously with purified GPX4 to form the expected 434 Da GPX4 adduct (Figure 6G). Together, these data are consistent with the protein reactivity of the proposed nitrile oxide electrophile, while simultaneously revealing surprising cellular selectivity in spite of this reactivity.

**Figure 6.**
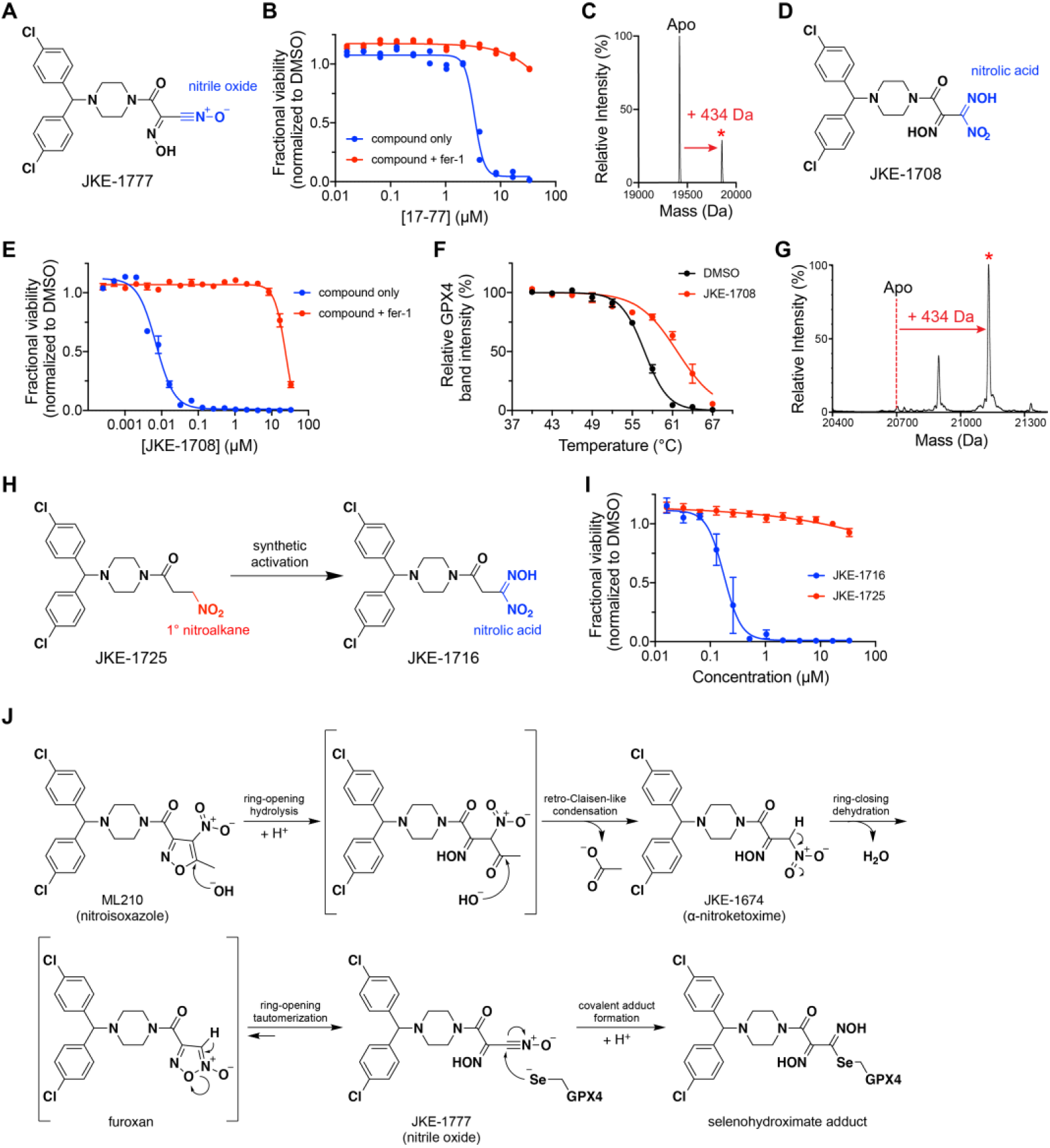
Dehydration of JKE-1674 yields a nitrile oxide electrophile that binds GPX4 covalently. (A) Proposed structure of JKE-1777 with nitrile oxide group shown in blue. (B) Co-treatment with fer-1 rescues the cell-killing effects of JKE-1777 in LOX-IMVI cells. Data are plotted as two individual technical replicates. (c) JKE-1777 is able to form a covalent adduct with purified GPX4^U46C^ protein. Adduct peak is marked with a red asterisk (*). (D) Structure of nitrolic acid JKE-1708 with nitrolic acid group shown in blue. (E) Co-treatment with fer-1 rescues the cell-killing effects of JKE-1708 in LOX-IMVI cells. Data are plotted as mean ± s.e.m., n = 4 technical replicates. (F) GPX4 CETSA of intact cells treated with JKE-1708 (red) reveals thermal stabilization of GPX4 compared to treatment with DMSO (black). Data are plotted as mean ± s.e.m., n ≥ 3 biologically independent samples. (G) JKE-1708 forms covalent adduct when incubated with purified GPX4 protein. Adduct peak is marked with a red asterisk (*). (H) Inactive compound JKE-1725, containing a primary nitro group, is activated upon synthetic transformation to nitrolic acid JKE-1716. (I)Viability measurements of cells treated with inactive nitroalkane JKE-1725 (red) and active nitrolic acid JKE-1716 (blue). Data are plotted as mean ± s.e.m., n = 4 technical replicates. (J) Proposed cellular activation pathway of masked nitrile oxide GPX4 inhibitors.

Finally, the discovery of JKE-1777 and JKE-1708 enabled determination of the GPX4 adduct structure formed by ML210-derived compounds. This was achieved by reacting JKE-1777 and JKE-1708 with small-molecule thiols (Figure S7D). Both JKE-1777 and JKE-1708 generate a thiol adduct containing a thiohydroximate group (Figure S7E). Similar thiohydroximate adducts of cysteine and glutathione also form in cells treated with ML210, JKE-1674, or JKE-1708 (Figures S5D and S5E). Based on these observations, ML210-derived inhibitors likely bind GPX4 to form the analogous selenium-containing structure.

### Proposed cellular activation pathway of masked nitrile oxide GPX4 inhibitors

Our findings reveal that cellular conversion of ML210 into its active nitrile oxide form occurs through a unique two-step sequence involving hydrolysis of the nitroisoxazole (Figure 5C) and dehydration of the resulting α-nitroketoxime (Figure S6A). We were unable to directly dehydrate α-nitroketoxime JKE-1674 and instead relied on a reduction-oxidation sequence to prepare nitrile oxide JKE-1777 via an intermediate bis-oxime (Figure S6B). This intermediate does not induce ferroptosis (Figure S8A), suggesting that the dehydration of JKE-1674 in cells proceeds through a different pathway.

To gain further insight into the process by which an electrophile is produced from the dehydration of JKE-1674 in cells, we performed SAR studies focused on the α-nitroketoxime group (Figure S8A). Like the ML210 nitroisoxazole, this α-nitroketoxime moiety exhibits steep SAR (Figure S8A), indicating that both the oxime and nitro groups play crucial roles in enabling GPX4 binding. We found that inactive JKE-1674 analogs containing a primary nitroalkane but lacking an oxime can be synthetically converted into cell-active GPX4 inhibitors by incorporation of a nitrolic acid group (Figures 6H, 6I, and S8C). These observations indicate that the oxime group is not a vestigial structure of the ML210 nitroisoxazole ring, but plays a critical role in the cellular conversion of the α-nitroketoxime in JKE-1674 to an electrophile. A possible explanation is that dehydration of the α-nitroketoxime initially yields an unstable monosubstituted furoxan heterocycle (Figure 6J), the formation of which is dependent on the presence of the oxime group. The furoxan, in turn, can readily undergo a ring-opening isomerization reaction to yield a nitrile oxide^21,22^. While either this furoxan or the isomeric nitrile oxide JKE-1777 could plausibly act as an electrophile, other nitrile oxide precursors that are incapable of forming furoxans are also able to engage GPX4 (Figures 6H, 6I, and S8C). This observation is consistent with a nitrile oxide as the most likely ML210-derived electrophile (Figure 6J).

### Pharmacokinetic advantages of masked electrophiles

Our efforts to illuminate the mechanism of action of ML210 have identified a family of previously uncharacterized masked electrophiles that, upon cellular activation, covalently inhibit GPX4. The masked reactivity of these molecules promises to overcome key pharmacokinetic liabilities of existing GPX4 inhibitors, which rely on highly-reactive chloroacetamide (or chloroacetamide-like) warheads to covalently inhibit GPX4^4,5^. Consistent with the expected shortcomings of highly reactive covalent inhibitors, we experimentally observed that chloroacetamides RSL3 and ML162 do indeed exhibit poor physiochemical and pharmacokinetic properties (Table 1). Attempts to improve upon these properties by using alternate warheads with attenuated electrophilic groups, such as clinically relevant acrylamides, produced inactive analogs (Figures S9A and S9B).

**Table 1.** Physiochemical and pharmacokinetic properties of GPX4 inhibitors.

| Compound |  | ML210 | ML162 | RSL3 | JKE-1674 |
| --- | --- | --- | --- | --- | --- |
| MW (g/mol) |  | 475.3 | 477.4 | 440.9 | 451.3 |
| cLogP |  | 4.7 | 4.8 | 3.4 | 4.0 |
| logD (pH 7.5) |  | 4.9 | 3.6 | 3.0 | 3.1 |
| Solubility (mg/L) |  | <0.1 | 0.2 | * | 45.6 |
| Caco A-B [nm/s] |  | * | * | * | 113.2 |
| Efflux ratio |  | * | * | * | 1.12 |
| Plasma stability (%) | human | 21 | 1 | 27 | 100 |
|  | rat | 76 | 21 | 0 | 100 |
|  | mouse | 24 | 27 | 0 | 100 |
| Buffer stability (%) | pH 1 | 79 | 100 | 87 | 100 |
|  | pH 7 | 100 | n.d. | n.d. | 100 |
|  | pH 10 | 0 | 100 | 39 | 100 |
| Glutathione stability (%) |  | 69 | 100 | 100 | n.d. |
| Cysteine stability (%) |  | 26 | 0 | 0 | n.d. |
| Rat hepatocyte stability (Fmax) (%) |  | 20 | <1 | <1 | 76 |
| Microsome stability (Fmax) (%) | human | 47 | 9 | 7 | 34 |
|  | mouse | 41 | <1 | 33 | 91 |

\* Value cannot be reported due to low compound recovery; n.d., not determined. cLogP, calculated log(partition-coefficient); LogD, log(distribution-coefficient) determined at pH 7.5.

The discovery and development of masked electrophiles represents an alternative approach to realize covalent inhibitors with improved physiochemical and pharmacokinetic properties. Masked electrophiles are inert to covalent protein binding until they are chemically transformed into active electrophilic species through enzymatic or non-enzymatic processes. As a result, masked electrophiles can be administered to cells or animals in a stable, non-reactive form and can enable selective targeting of cellular proteins by limiting exposure of the protein-reactive species to the site of action^25^. Consistent with these concepts, both ML210 and JKE-1674 exhibit far greater stability than chloroacetamide inhibitors (Table 1). In addition, the α-nitroketoxime group of JKE-1674 significantly improves upon the solubility of ML210 and chloroacetamide inhibitors (Table 1), crossing a critical threshold necessary for formulation and dosing in animals. Indeed, even without medicinal chemistry optimization, JKE-1674 can be detected in the serum of mice dosed orally with a PEG400/ethanol solution of the compound (Figure S9C). It is likely that systematic optimization of JKE-1674 and related masked nitrile oxides will yield further-improved tool compounds for understanding the basic biology of GPX4 and therapeutic potential of ferroptosis induction.

### Use of ML210 and derivatives as tool compounds for elucidating GPX4 biology

To encourage experimental adoption of key cellular probes of GPX4 representing three different chemotypes (chloroacetamide, nitroisoxazole, α-nitroketoxime) (Figure 7A), we now detail their use in cellular assays relevant to ferroptosis. We confirm that all three chemotypes, as well as genetic loss of GPX4^26^, share a pattern of cell killing across a range of cancer cell lines (Figure 7B). A key distinction between these chemotypes concerns the degree to which their viability effects can be rescued by ferroptosis inhibitors fer-1 and zileuton, which have been shown to prevent lipid peroxidation by acting as radical-trapping antioxidants^12,27^. We found that the viability effects of masked nitrile oxide GPX4 inhibitors are rescued by ferroptosis inhibitors to a far greater degree than that of chloroacetamide GPX4 inhibitors (Figures 7C). This greater selectivity of masked nitrile oxide GPX4 inhibitors affords a number of additional advantages for their use in biological assays. For example, the loss of genes (e.g. *ZEB1*) that are necessary for ferroptotic death can rescue cells from death induced by all concentrations of ML210 (Figure 7D), without evidence of off-target, non-rescuable effects at high concentrations (>10 μM)^1^. As a result, when performing rescue experiments systematically, for example in a genome-scale CRISPR screen to illuminate the cellular circuitry regulating ferroptosis^1^, a fixed high concentration of ML210 (e.g.10 μM) can be chosen and used with convenience. In contrast, chloroacetamides must be titrated carefully to balance GPX4 inhibition against off-target effects and to facilitate validation and reproduction of results by others.

**Figure 7.**
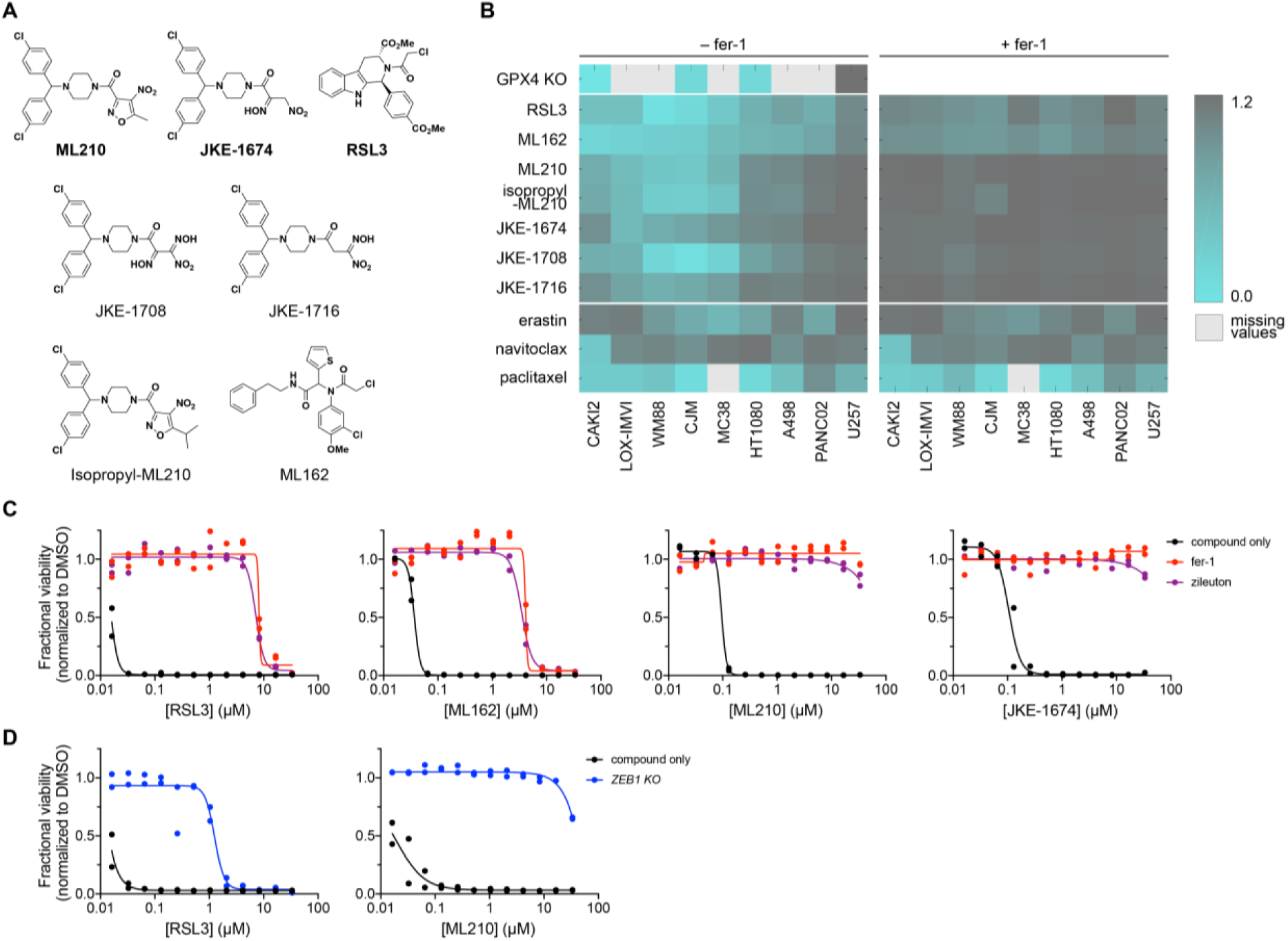
Profiling of structurally diverse GPX4 inhibitors in ferroptosis assays. (A) Chemical structures of diverse GPX4 inhibitors. Key tool compounds for cellular assays are bolded. (B) GPX4 inhibitors bearing chloroacetamide, nitroisoxazole, and α-nitroketoxime warheads exhibit a similar pattern of cell killing across a range of cancer cell lines relative to control lethal agents. Nitrile oxide precursors show enhanced rescuability by fer-1 relative to chloroacetamide GPX4 inhibitors. WM88, LOX-IMVI, CJM and U257 are human melanoma cell lines. CAKI2 and A498 are human renal cell carcinoma cell lines. HT1080 is a human fibrosarcoma cell line. MC38 is a mouse colon cancer cell line and PANC02 is a mouse pancreatic cancer cell line. (C) ML210 and JKE-1674 exhibit fewer off-target effects that cannot be rescued by radical-trapping antioxidants (e.g. fer-1, zileuton) in LOX-IMVI cells compared to chloroacetamide inhibitors. Data are plotted as two individual technical replicates. (D) Deletion of *ZEB1*, which is required for ferroptosis in KP4 pancreatic cancer cells, confers markedly greater rescue of cell killing by ML210 compared to RSL3. Data are plotted as two individual technical replicates.

Alongside a discussion of the advantages of masked nitrile oxide GPX4 inhibitors in biological experiments, we also note several caveats. The first relates to diminished potency of this class of inhibitors in certain contexts relative to chloroacetamide GPX4 inhibitors (Figure 7B). The second concerns potential discrepancies in the activity of masked nitrile oxide versus chloroacetamide GPX4 inhibitors in one out of two mouse cancer cell lines that we have examined (Figure 7B), an observation that warrants further investigation. This and other yet-to-be-discovered nuances between different GPX4-inhibiting chemotypes argues for the inclusion of several chemotypes in all biological experiments probing ferroptosis. A final caution relates to the need for careful storage and handling of JKE-1674, which can decompose if stored for prolonged periods at room temperature as a solution in DMSO.

## DISCUSSION

Drugs that form covalent bonds with their targets (covalent inhibitors) constitute important medicines in pharmacopeia (e.g. penicillin, aspirin, omeprazole, afatinib, ibrutinib) but historically have raised concerns about potential lack of selectivity. Recent work has renewed interest in the power of covalent binders for expanding the druggable proteome and is reversing dogma around the use of electrophilic small molecules in drug development^28–32^. In particular, several seminal studies have shown that a significant proportion of the undruggable proteome contains chemically accessible nucleophilic residues that can be exploited using chloroacetamide groups^28,31,32^.

Yet, challenges remain. Unfortunately, as we and others^33^ have shown, chloroacetamide-based tools are poor starting points for further development given their promiscuity, instability, and low bioavailability. Therefore, therapeutic translation of the many tantalizing leads generated by recent ligandability studies^28,31,32,34–36^ requires substitution of the chloroacetamide warhead with a more drug-like electrophilic group. While acrylamides or other attenuated electrophiles may be appropriate alternatives in certain cases^33^, in other contexts, they may be insufficient to substitute for more reactive warheads, such as chloroacetamides^31^.

GPX4 is a protein that exemplifies this challenge. While GPX4 is readily modified by chloroacetamide-containing compounds, our extensive efforts to replace this functional group with acrylamides or other protein-reactive electrophiles have revealed that conventional alternatives are not suitable for targeting GPX4. This trend is corroborated by the abundance of chloroacetamide-containing hits and absence of acrylamide-containing hits in high-throughput screens aiming to identify GPX4 inhibitors^6,37,38^. Related results suggest that GPX4 is unlikely to be unique in this regard among the hundreds of proteins that are accessible through chloroacetamide-containing tool compounds^31^. For example, a screen to identify inhibitors of GSTO1 enriched highly for chloroacetamide hits but not those containing acrylamides or other drug-like warheads^39^. For such proteins, it is challenging to identify drug-like warheads that can substitute for the reactivity of chloroacetamides^40^ and precedent for the development of drug-like inhibitors is absent^33^.

In this manuscript, we describe our discovery that nitrile oxide electrophiles comprise an effective strategy to substitute for chloroacetamide groups within GPX4 inhibitors. GPX4 inhibitors that act via nitrile oxides exhibit remarkable selectivity for GPX4 and diminished off-target effects compared to chloroacetamide-based GPX4 inhibitors. The discovery that nitrile oxide electrophiles can engage in such highly selective cellular interactions is unexpected given that nitrile oxides are regarded as one of the most reactive organic functional groups^41^. Indeed, the reactivity of nitrile oxides has previously precluded their isolation and investigation as protein-reactive electrophiles^42,43^. The biological utility of nitrile oxide electrophiles that we have now uncovered is predicated on the unique electrophile-masking elements present in the ML210 nitroisoxazole chemical pathway. These prodrug features, in addition to enabling application to cells and mechanistic interrogation, also bestow masked nitrile oxides with pharmacokinetic properties and bioavailability that far surpass those of unmasked electrophiles (e.g. chloroacetamides).

Intriguingly, the cellular selectivity of masked nitrile oxides does not appear to arise from their concealed reactivity. Instead, our results suggest that cellular selectivity may, unexpectedly, be a feature intrinsic to nitrile oxides themselves. For example, the unmasked nitrile oxide JKE-1777, which can be isolated by virtue of oxime stabilization of the nitrile oxide^21^, exhibits selectivity equivalent to masked precursors as assessed by multiple orthogonal assays. We speculate that it may be the transient nature of nitrile oxides that prevents prolonged exposure of cells to this highly reactive electrophilic species and thereby avoids extensive covalent interactions with the proteome. While our work has been concerned primarily with the relevance of masked nitrile oxides for the development of GPX4 inhibitors, the general properties of masked nitrile oxides that we describe are likely to be extensible. For example, proof-of-concept data using a structurally distinct masked nitrile oxide suggest that this functionality may be a useful warhead for achieving selective, covalent interactions with cellular proteins beyond GPX4 (Figure S10).

The toolkit of GPX4-inhibiting compounds uncovered by our mechanistic studies represents an unprecedented wealth of diverse GPX4-targeting chemotypes (nitroisoxazoles, α-nitroketoximes, and nitrolic acids). These compounds promise to overcome a long-standing shortcoming of the GPX4-biology field, namely, reliance on a single non-specific chemotype (chloroacetamides) to perform and interpret biological experiments^4,44^. This liability is the chemical biology equivalent of drawing conclusions from functional genetic experiments performed with a single short hairpin or guide RNA^45–47^. We therefore encourage scientists probing GPX4 and ferroptosis biology to incorporate into their experimental designs the chemically diverse GPX4 inhibitors we have illuminated. This approach promises to mitigate chemotype-specific off-target effects and facilitate the generation of robust, reproducible, and extensible results^45–48^.

Finally, we expect that the numerous tool compounds, mechanistic insights, cellular target engagement assays, structural information, and biochemical data we make available will be catalytic for the development of GPX4-inhibiting drugs by the academic and pharmaceutical communities.

## SUPPLEMENTAL INFORMATION

## ACKNOWLEDGMENTS

The authors would like to thank Virendar Kaushik for assistance with intact protein mass spectrometry experiments; Michael J. Palte for assistance with preparing phosphatidylcholine hydroperoxide; Bogdan A. Budnik and Renee A. Robinson for assistance with proteomics; Marion Slezewski for assistance with protein crystallization; Jutta Hoffmann and Lennart Schnirch for MS time course experiments to identify conditions for co-crystallization and for crystallographic refinement work; William Ho and Suvruta Iruvanti for helpful comments on this manuscript. This work was supported by grants from the National Institutes of Health (R01GM038627 and R35GM127045) to S.L.S. It was also supported in part through a collaboration between the Broad Institute and Bayer AG. D.M., R.C.H., A.H., M.N., V.B., S.G., S.Ch., R.N., and A.L.E. are employed by Bayer AG.

## AUTHOR CONTRIBUTIONS

J.K.E. conceived of, designed, and performed experiments. L.F., K.E.L., S.G. and C.M. performed chemical synthesis. A.L.E. performed *in vivo* experiments. R.A.R., M.J.R. and V.S.V. maintained cell cultures and performed viability, cellular thermal shift, and western blotting experiments. A.H. and D.M. designed the cloning approach and expressed, purified, characterized recombinant wild type GPX4 protein, and performed the in-cell compound-binding assay. R.C.H. designed and supervised the protein crystallization and structure determination study. V.B. performed intact protein mass spectrometry studies. M.N. performed metabolite-ID studies. A.K., S.C., and B.B. contributed tools and reagents for protein characterization experiments. P.A.C. contributed to analysis of cell viability data. S.Ch. performed cellular lipid peroxidation assays. R.N. performed formulation work. J.K.E., V.S.V. and S.L.S. initiated the project and wrote the manuscript. V.S.V. and S.L.S. directed the project.

## EXTENDED DISCUSSION

Biochemical and structural studies of GPX4 and its inhibitors have been hindered by the lack of an efficient source of wild-type GPX4 protein^5,49,50^. The recombinant expression of mammalian selenoproteins in bacterial protein production systems is often difficult due to low efficiency of selenocysteine incorporation^51–53^. Historically, structural studies of GPX4 have been possible only with U46C and U46G active-site mutants^38,52,54,55^. The structure of a selenocysteine-containing GPX4 (though not wild type) was recently reported^49^, but to date, structural studies of covalent GPX4 inhibitors have not been successful^5^. Co-crystal structures of GPX4^U46C^ in complex with reversible peptide binders have been reported, but their relevance remains unknown given that these compounds are unable to inhibit cellular GPX4^38^.

In this manuscript, we report the first crystallization and structural determination of true wild-type GPX4 protein as well as of the first complex of GPX4 with a covalent small-molecule inhibitor. This was achieved by co-expression of GPX4 with SECIS-binding protein 2 (SBP2) in HEK293-6E cells^51^, which allowed generation of true wild-type GPX4 and GPX4^C66S^ proteins. The apo wild-type GPX4 structure described here is similar to mutant GPX4 structures that have been previously reported. However, it does not show any signs of oxidation on the selenium atom of U46, unlike what has been reported recently for a U46-containing GPX4 protein produced via a different strategy^49^. This difference in oxidation may be related to the 10-fold higher concentration of the reducing reagent TCEP used in our study (see Methods).

Generation of GPX4 crystals in complex with a GPX4 inhibitor (ML162) was initially complicated by heterogeneous modification of GPX4 by ML162, including partial covalent modification of the surface-exposed C66 residue at high concentrations of ML162. Mutation of C66 to serine, along with optimization of binding reaction conditions and use of enantiomerically-pure (*S*)-ML162, enabled determination of a co-crystal structure at 1.54 Å resolution. ML162 is fully defined in the electron density and covalently linked to U46 (Figure 4A and Table S1, data set GPX4:ML162(S)). The amide moiety adjacent to the covalent bond to U46 is present in two alternative conformations where the oxygen atom forms a hydrogen bond to the side chains of Trp136 and of Asn137, respectively. The oxygen of the second ML162 amide group is coordinated via a hydrogen bond to the backbone nitrogen of Gly47, and the methoxyphenyl moiety is in π-π stacking distance to the side chain of Gln 81.

Our results provide the first insight into the structural basis of GPX4 inhibition by chloroacetamide inhibitors. The lack of a well-defined binding pocket in the ML162 co-crystal structure reaffirms the difficulties associated with targeting GPX4 with small-molecules. Based on these observations, it is unsurprising that currently known inhibitors target GPX4 through a covalent mechanism of action. Using the GPX4 expression and co-crystallization strategy that we have described may enable the structural characterization of other GPX4 binders, which could guide rational development efforts of improved inhibitors.

## SUPPLEMENTARY FIGURES

**Figure S1.**
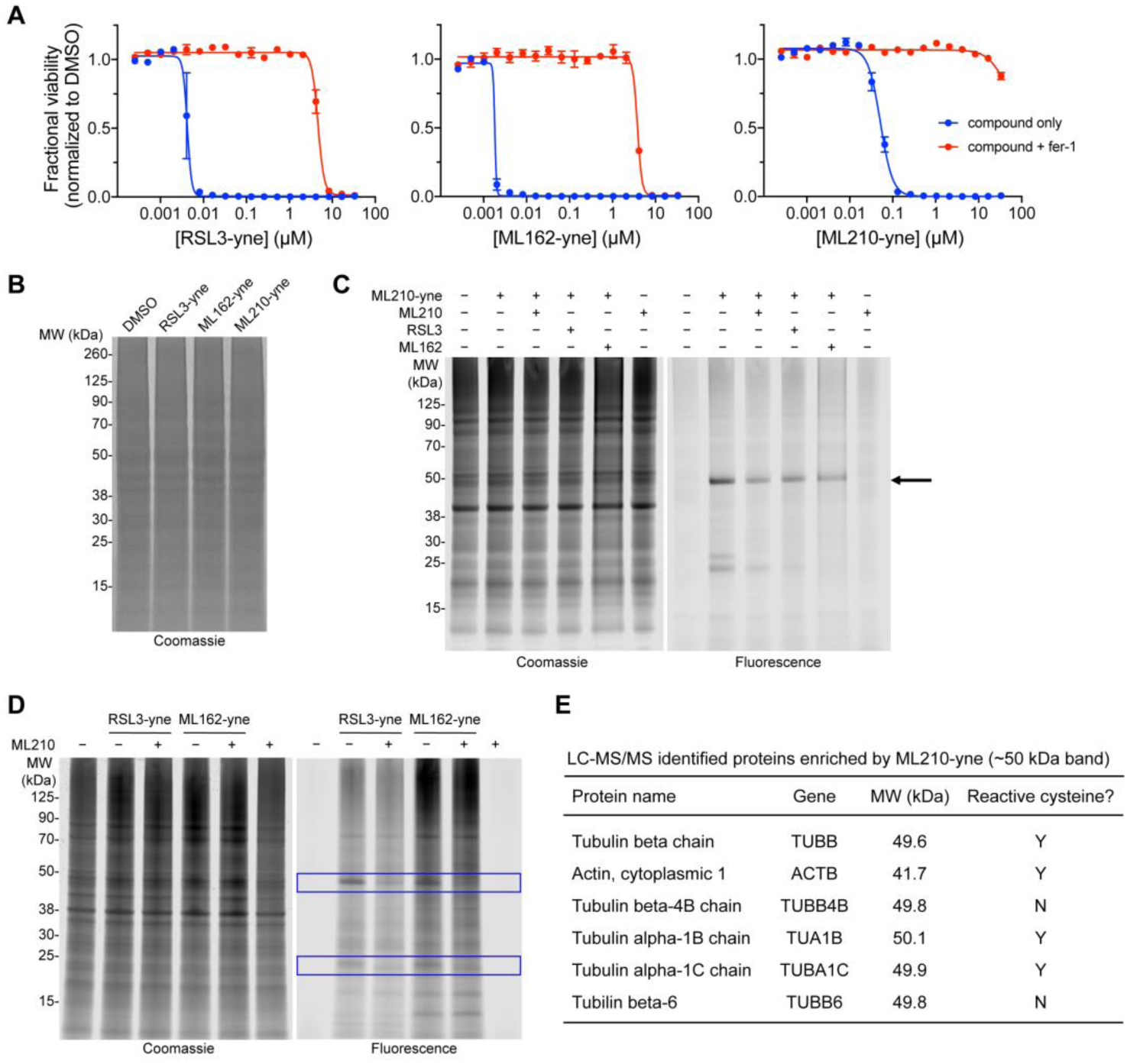
Characterization of GPX4-inhibitor alkyne probe analogs, related to Figure 2. (A) Alkyne analogs RSL3-yne, ML162-yne, and ML210-yne exhibit cellular activity similar to their respective parent compounds. Data are plotted as mean ± s.e.m., n = 4 technical replicates. (B) Coomassie-stained gel corresponding to fluorescence gel in main text Figure 3A. (C) (Competitive cellular reactivity profile of ML210-yne. Arrow indicates ≄50 kDa band, which is not a specific target of ML210-yne. (D) Competitive cellular reactivity profiles of RSL3-yne and ML162-yne. Blue boxes indicate bands that are shared targets of ML210 and chloroacetamide probes. (E) Table of proteins enriched by ML210-yne in ≄50 kDa band. The most highly enriched proteins are abundant cytoskeletal proteins, many of which contain previously characterized reactive cysteine residues^32,56^.

**Figure S2.**
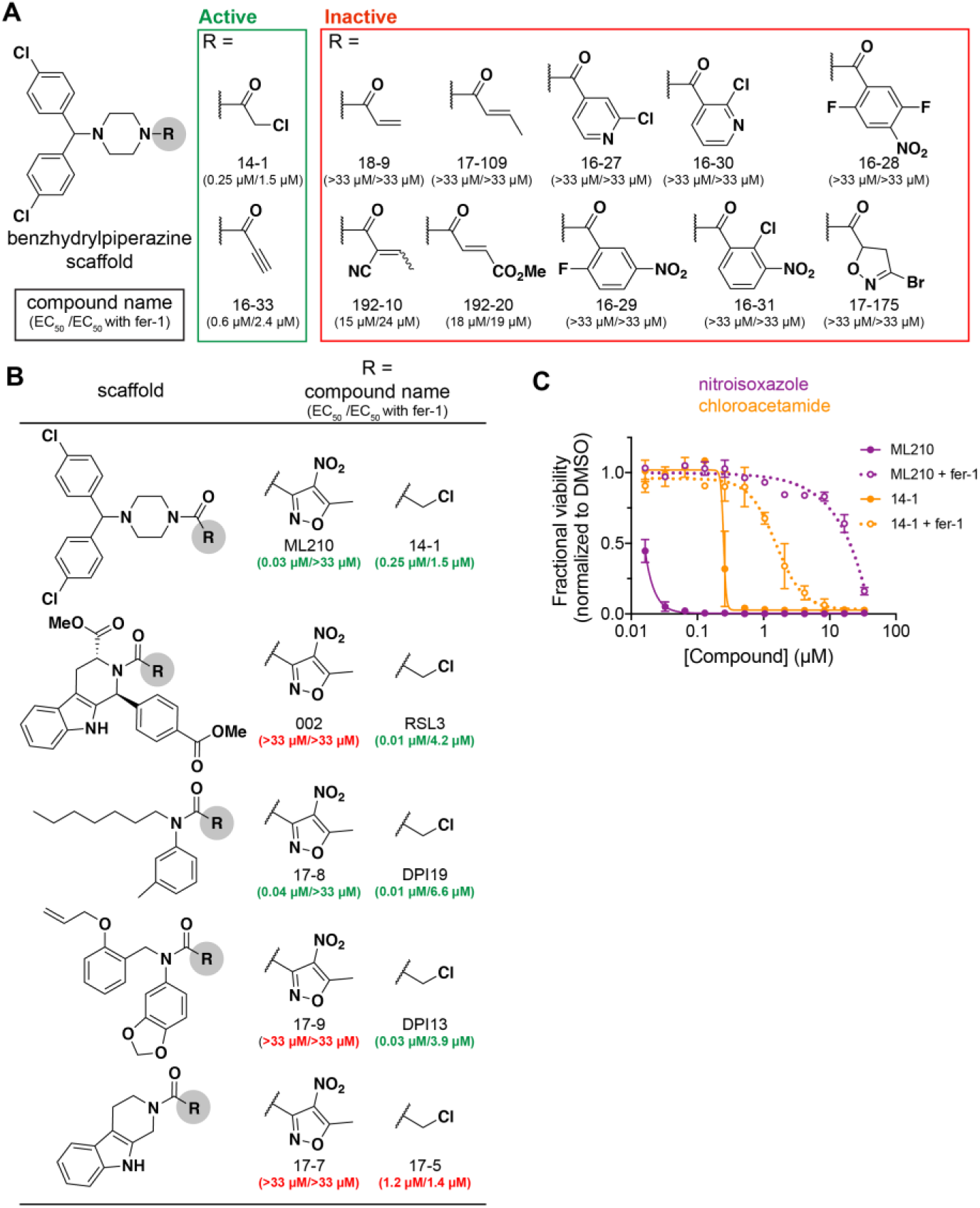
Assessment of GPX4-targeting electrophilic warheads, related to Figure 2. (A)Replacement of the nitroisoxazole warhead of ML210 with other electrophilic groups reveals chloroacetamide and propiolamide analogs are far less selective for GPX4. EC_50_ values were determined in LOX-IMVI cells from 12-point dose-response experiments (n ≥ 2 technical replicates). (B) Replacement of the chloroacetamide warhead of GPX4 inhibitors with the nitroisoxazole warhead of ML210 can enhance selectivity. For some chloroacetamide GPX4 inhibitors, including RSL3, this substitution with nitroisoxazole is not tolerated. EC _50_ values were determined in LOX-IMVI cells from 12-point dose-response experiments (n ≥ 2 technical replicates). (C) ML210 is more potent and selective than the corresponding analog with the nitroisoxazole warhead replaced with a chloroacetamide group. Data are from measurements of LOX-IMVI viability and are plotted as mean ± s.e.m., n = 4 technical replicates.

**Figure S3.**
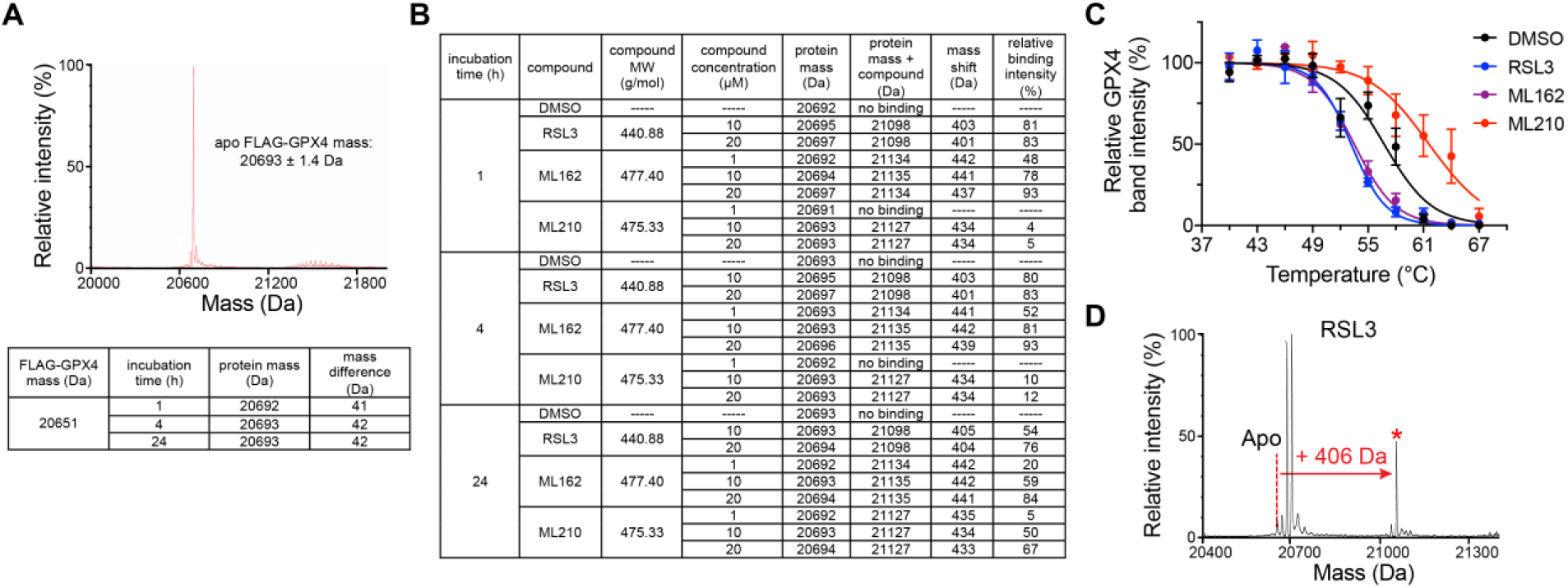
Characterization of GPX4 inhibitor binding, related to Figures 3 and 4. (A) Characterization of in-cell FLAG-GPX4 binding by intact protein mass spectrometry. Upper plot depicts deconvoluted apo FLAG-GPX4 mass spectrum. Table summarizes proteins masses detected after incubation of cells for 1, 4, or 24 h. Observed protein mass indicates that FLAG-GPX4 is acetylated in cells. Standard deviation of apo protein mass was determined from n = 48 technical replicates. (B) Summary of in-cell binding experiments. Compounds (1, 10, or 20 μM) were incubated with cells expressing FLAG-GPX4 for 1, 4, or 24 hours. Observed RSL3 and ML162 adduct masses are consistent with loss of HCl (36 Da) upon reaction with GPX4. ML210 binding is associated with a loss of 41 Da. (C) GPX4 CETSA profiles for intact cells (HT1080) treated with DMSO (black), RSL3 (blue), ML210 (red), or ML162 (purple). Cells were treated with 10 μM compound for 1 h. Data are plotted as mean ± s.e.m., n ≥ 2 biologically independent samples. (D) RSL3 binds GPX4^U46C^ in cells.

**Table S1.** Crystallographic data collection and refinement statistics, related to Figure 4.

| Data set | GPX4 (apo) | GPX4:ML162(racemate) | GPX4:ML162(S) |
| --- | --- | --- | --- |
| Space group | *P*1 | *P*2<sub>1</sub> | *P*2<sub>1</sub>2<sub>1</sub>2<sub>1</sub> |
| Cell dimensions |  |  |  |
| *a*, *b*, *c* (Å) | 32.8, 35.2, 37.8 | 32.8, 68.4, 69.9 | 32.7, 57.2, 81.3 |
| α, β, γ (°) | 103.2, 112.3, 91.8 | 90.0, 92.5, 90.0 | 90.0, 90.0, 90.0 |
| Resolution (Å)* | 1.01–33.97<br>(1.01–1.03) | 2.29–48.88<br>(2.29–2.42) | 1.55–46.82<br>(1.55–1.59) |
| *R*<sub>sym</sub>* | 3.6 (52.3) | 19.3 (58.2) | 10.5 (90.2) |
| *I*/σ* | 14.6 (1.8) | 5.1 (2.0) | 20.2 (2.4) |
| Completeness (%)* | 94.7 (87.6) | 98.7 (96.4) | 86.6 (72.1) |
| Redundancy* | 1.8 (1.6) | 3.5 (3.3) | 10.5 (6.7) |
| Refinement |  |  |  |
| Resolution (Å) * | 1.01–33.97<br>(1.00–1.03) | 2.29–48.88<br>(2.29–2.35) | 1.54–46.81<br>(1.54–1.58) |
| No. reflections | 73153 | 13191 | 19002 |
| Rwork / Rfree | 11.49/13.40<br>(26.5/27.8) | 17.42/23.94<br>(23.3/28.9) | 14.53/18.93<br>(23.1/33.2) |
| No. atoms |  |  |  |
| GPX4 | 1471 | 2746 | 1365 |
| ML162 | n/a | n/a | 32 |
| Cl<sup>-</sup>/EtOH | 5 | n/a | n/a |
| SO<sub>4</sub>/Ethylenglycol/DMSO | n/a | n/a | 21 |
| Water | 254 | 200 | 230 |
| B-factors |  |  |  |
| GPX4 | 14.3 | 23.5 | 15.9 |
| ML162 | n/a | n/a | 25.2 |
| Cl<sup>-</sup>/EtOH | 18.2 | n/a | 39.8 |
| Water | 30.5 | 25.3 | 29.7 |
| R.m.s. deviations |  |  |  |
| Bond lengths (Å) | 0.011 | 0.018 | 0.020 |
| Bond angles (°) | 1.568 | 1.938 | 2.141 |

\*Values in parentheses are for highest-resolution shell.

**Figure S4.**
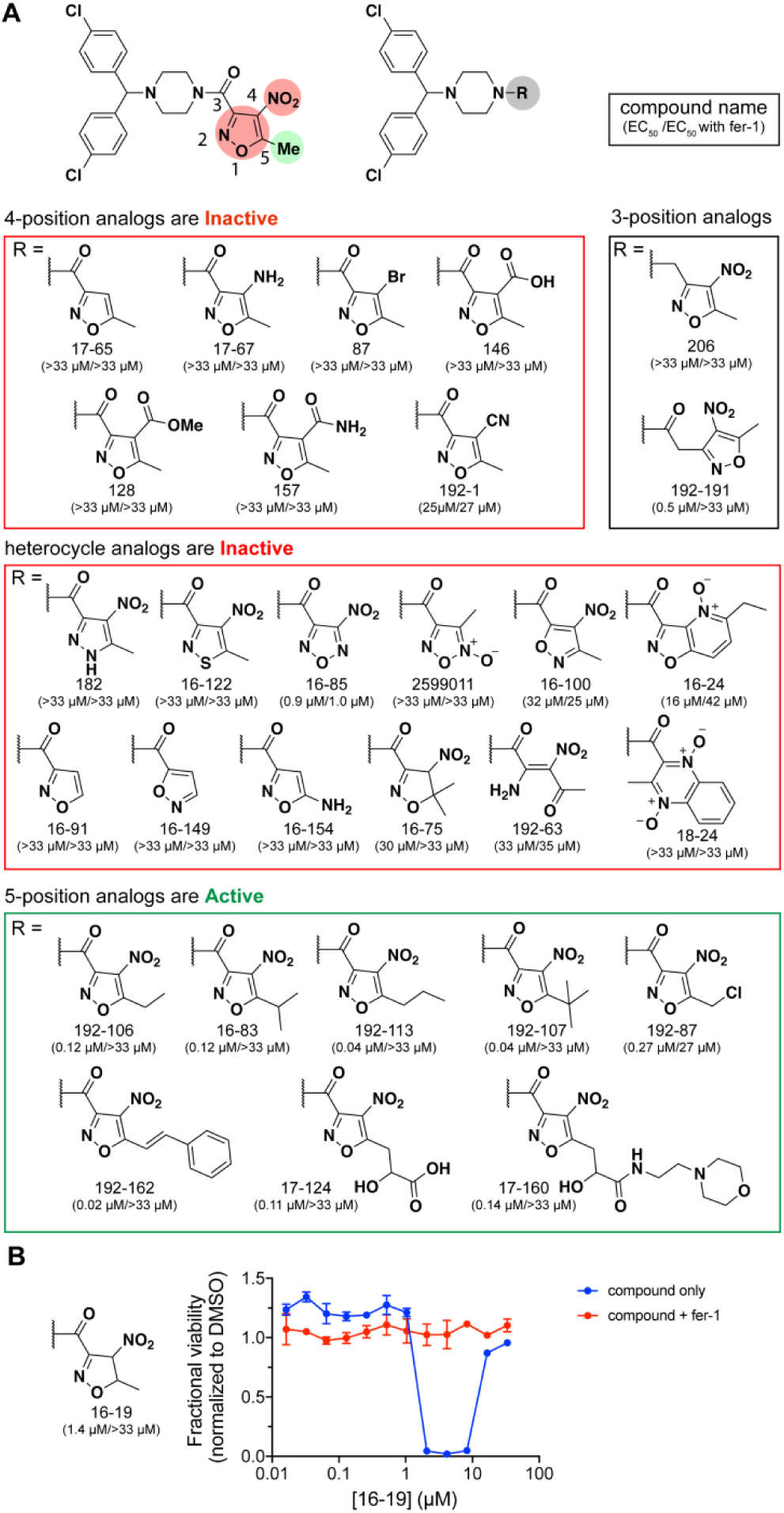
Summary of ML210 nitroisoxazole structure-activity relationship (SAR) study, related to Figure 5A. (A) ML210 analogs were synthesized and tested in cell viability assays in LOX-IMVI melanoma cells. Compounds are classified as inactive GPX4 inhibitors (red boxes) if they have no fer-1 rescuable effect on cell viability. The cellular effects of active analogs (green box) can be rescued by co-treatment fer-1. EC_50_ values were determined from 12-point dose-response experiments (n ≥ 2 technical replicates). (B) Dihydroisoxazole analog 16-19 exhibits activity rescuable by fer-1 and exhibits an unusual “bell-shaped” dose-response profile. Oxidation of the dihydroisoxazole group to a nitroisoxazole produces ML210, which may explain the cellular effects of 16-19. Data are plotted as mean ± s.e.m., n ≥ 2 technical replicates.

**Figure S5.**
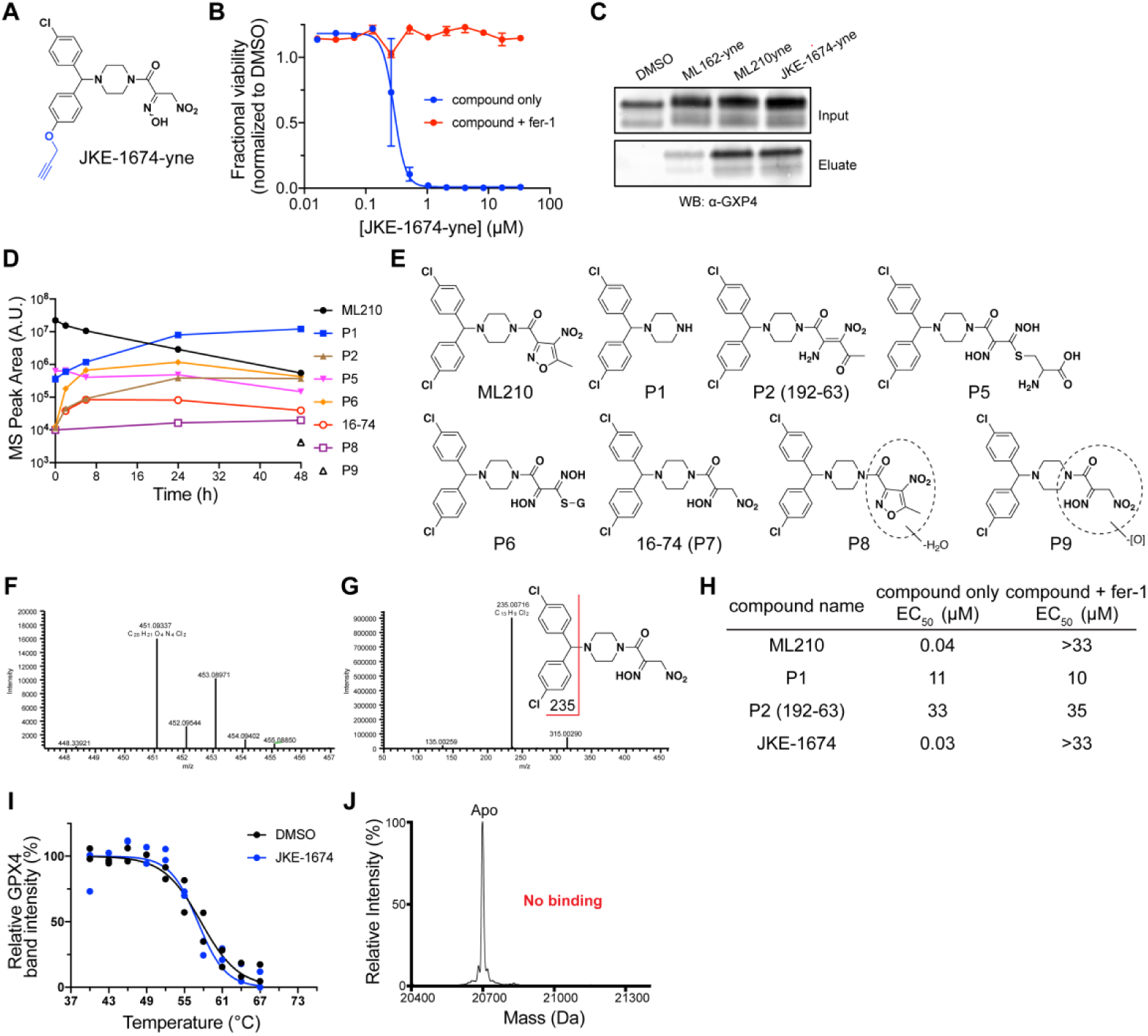
Cellular activity of ML210 and JKE-1674, related to Figure 5. (A) Structure of JKE-1674-yne. Alkyne group is highlighted in blue. (B) JKE-1674-yne exhibits cellular activity similar to JKE-16-74. Data are plotted as mean ± s.e.m., n ≥ 2 technical replicates. (C) JKE-1674-yne enables enrichment of GPX4 from cells to the same degree as ML210-yne. (D) Levels of ML210 and 7 related metabolites measured over the course of 48 hours in LOX-IMVI cells treated with ML210. Metabolites were measured by LC-MS/MS. (E) Chemical structures of ML210-derived metabolites. (F) MS spectrum of JKE-1674 observed in cells treated with ML210. These spectra are consistent with those of a synthetic standard of JKE-1674. (G) MS^2^ spectrum of JKE-1674 observed in cells treated with ML210. These spectra are consistent with those of a synthetic standard of JKE-1674. (H) Table of cell viability measurements in LOX-IMVI cells for selected metabolites. Only JKE-1674 exhibits cellular activity comparable to ML210. EC50 values were determined from 12-point dose-response experiments. (I) GPX4 CETSA profile for LOX-IMVI lysates treated with DMSO (black) or JKE-1674 (blue). Data are plotted as individual replicates from two separate biological experiments. (J) JKE-1674 does not interact with purified wild type GPX4 protein as determined by biochemical binding experiments using intact protein mass spectrometry.

**Figure S6.**
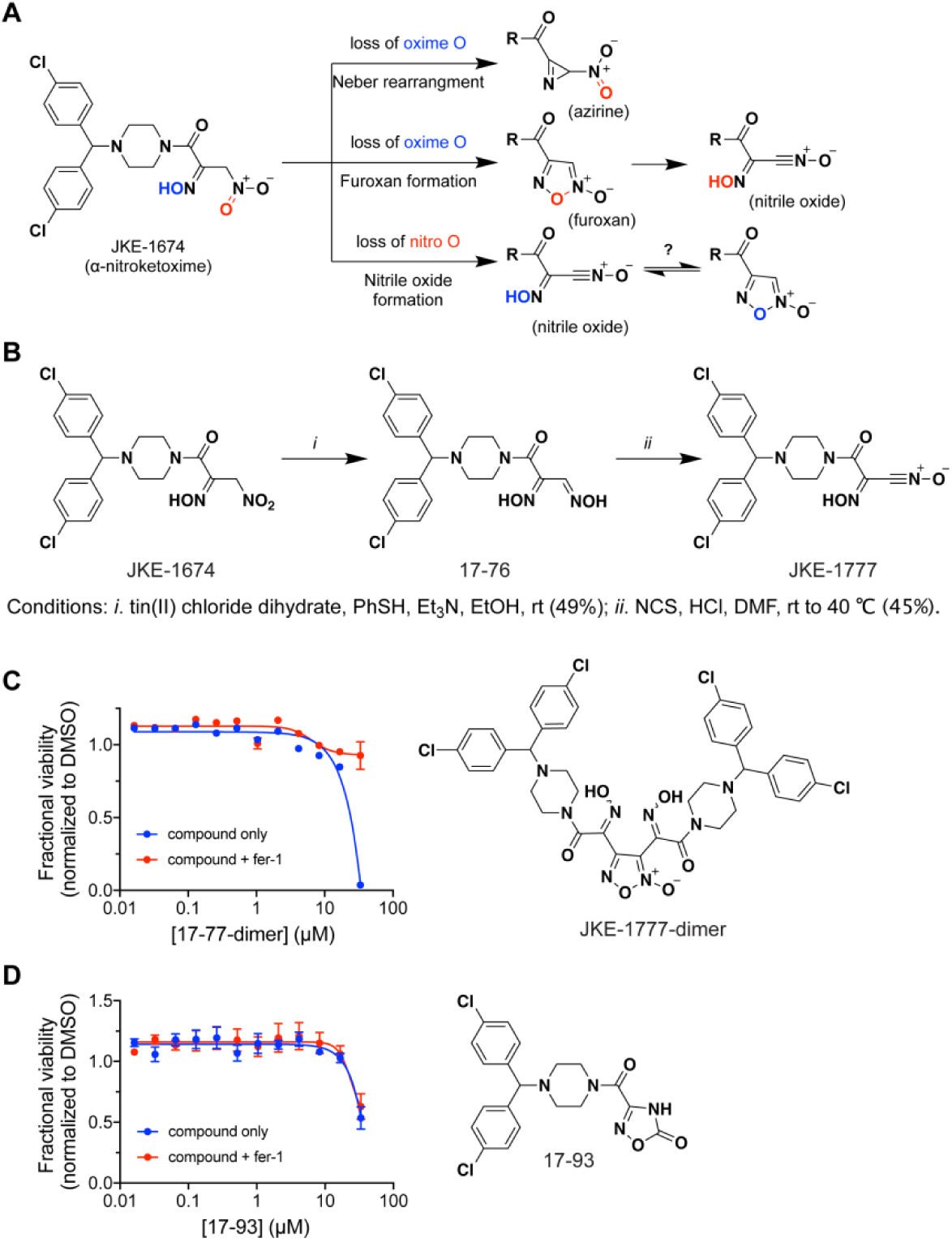
Identification of nitrile oxide JKE-1777, related to Figure 6. (A) Possible dehydration pathways of the JKE-1674 α-nitroketoxime group. (B) Two-step synthesis of JKE-1777 from JKE-1674. (C) Cell viability assessment in LOX-IMVI cells of inactive JKE-1777 dimerization product (JKE-1777-dimer) and proposed structure. Data are plotted as mean ± s.e.m., n ≥ 2 technical replicates. (D) Cell viability assessment in LOX-IMVI cells of the oxadiazolone-containing 17-93, which may be a nitrile oxide rearrangement product of JKE-1777. Data are plotted as mean ± s.e.m., n = 4 technical replicates.

**Figure S7.**
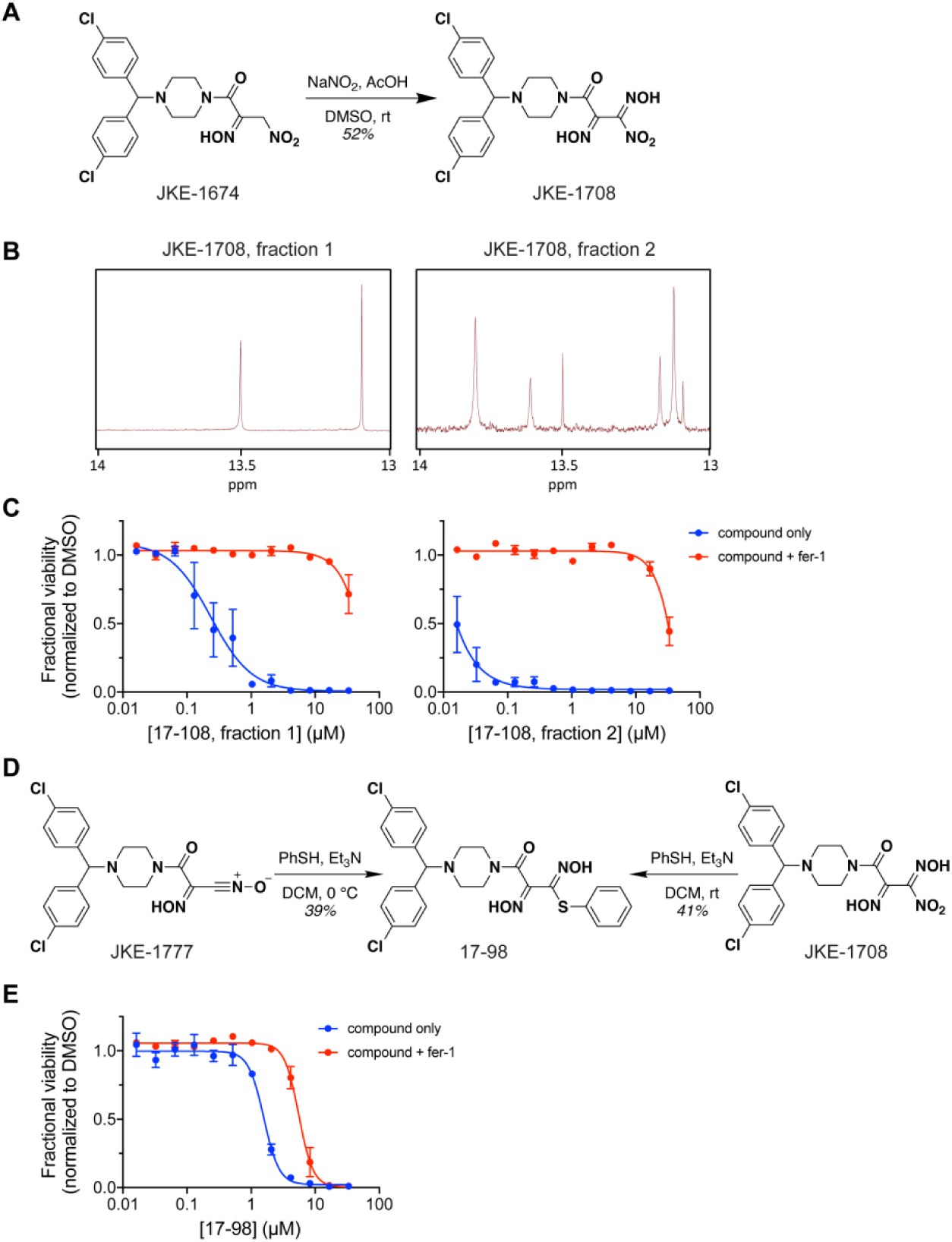
Characterization of nitrile oxide precursor GPX4 inhibitors, related to Figure 6. (A) Chemical synthesis of nitrolic acid JKE-1708 from JKE-1674. (B) ^1^H-NMR spectra of the –OH peaks (δ 13-14 ppm) of the two JKE-1708 fractions obtained after purification by flash column chromatography. Fraction 1 contains a single isomer while Fraction 2 contains 3 isomers. The ratio of isomers does not change for either compound in DMSO-*d6* for a period of at least 24 hours at room temperature. (C) JKE-1708 fraction 2 is more potent than JKE-1708 fraction 1. Data are plotted as mean ± s.e.m., n = 4 technical replicates. (D) Both nitrile oxide JKE-1777 and nitrolic acid JKE-1708 react with thiophenol to produce the thiohydroximate 17- 98. Both reactions produce slightly different mixtures of diastereomers; see Synthetic Methods. (E) Thiohydroximate 17-98 is a weak ferroptosis inducer and may act as a nitrile oxide precursor similar to JKE-1708. Data are plotted as mean ± s.e.m., n = 2 technical replicates.

**Figure S8.**
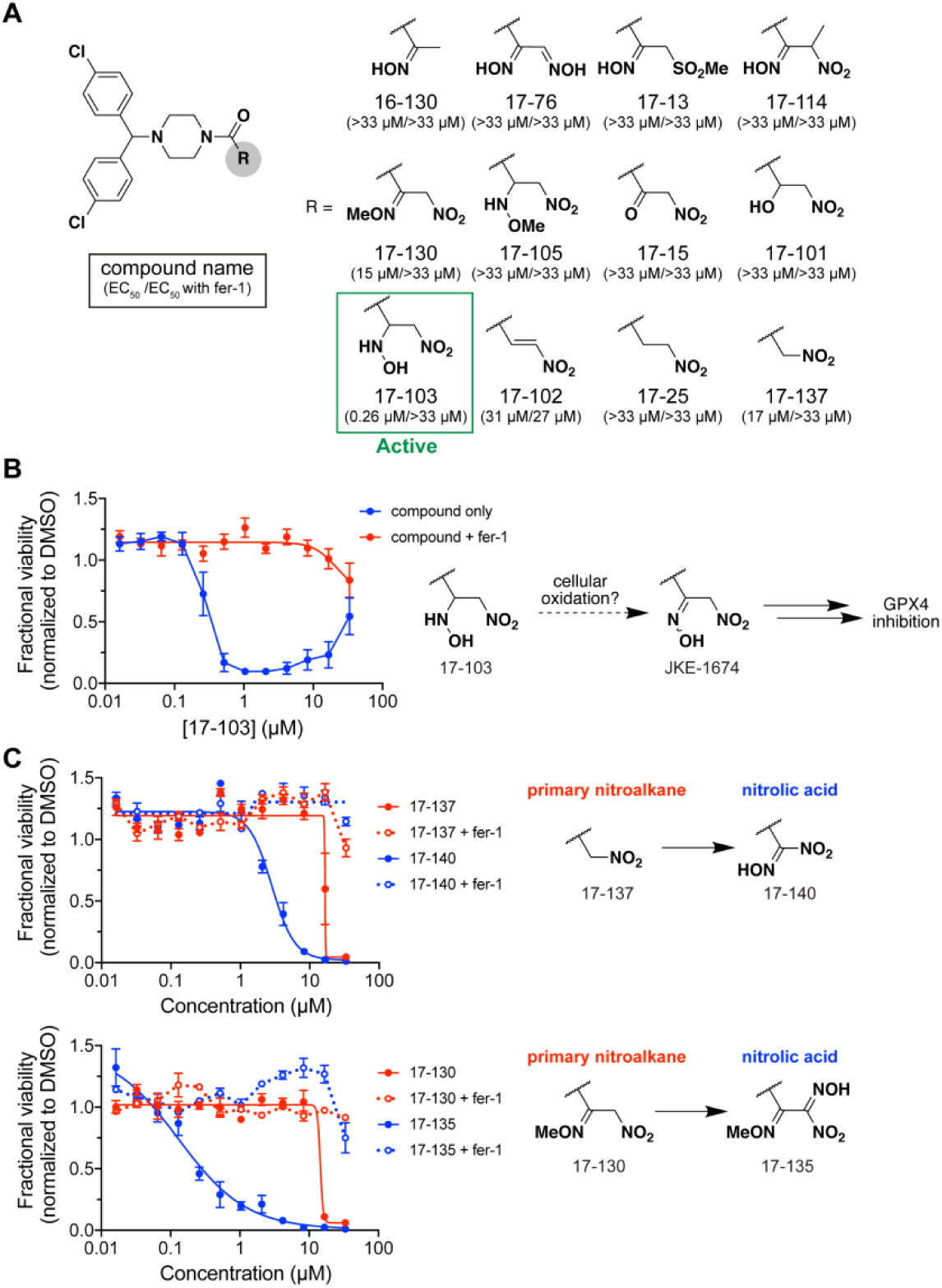
Structure-activity relationship (SAR) studies of JKE-1674, related to Figure 6. (A) Summary of JKE-1674 SAR. EC_50_ values were determined from 12-point dose-response experiments in LOX-IMVI cells (n ≥ 2 technical replicates). (B) Hydroxylamine 17-103 exhibits a ‘bell-shaped’ dose-response in cell viability assay. Oxidation of the hydroxylamine group to an oxime produces JKE-1674, which may explain the cellular effects of 17-103. Data are plotted as mean ± s.e.m., n = 6 technical replicates. (C) Primary nitroalkanes that do not effectively target GPX4 can be converted to active nitrolic acids. Fer-1 is able to rescue completely the cell-killing activity of nitrolic-acid GPX4 inhibitors. Data are plotted as mean ± s.e.m., n = 3 technical replicates.

**Figure S9.**
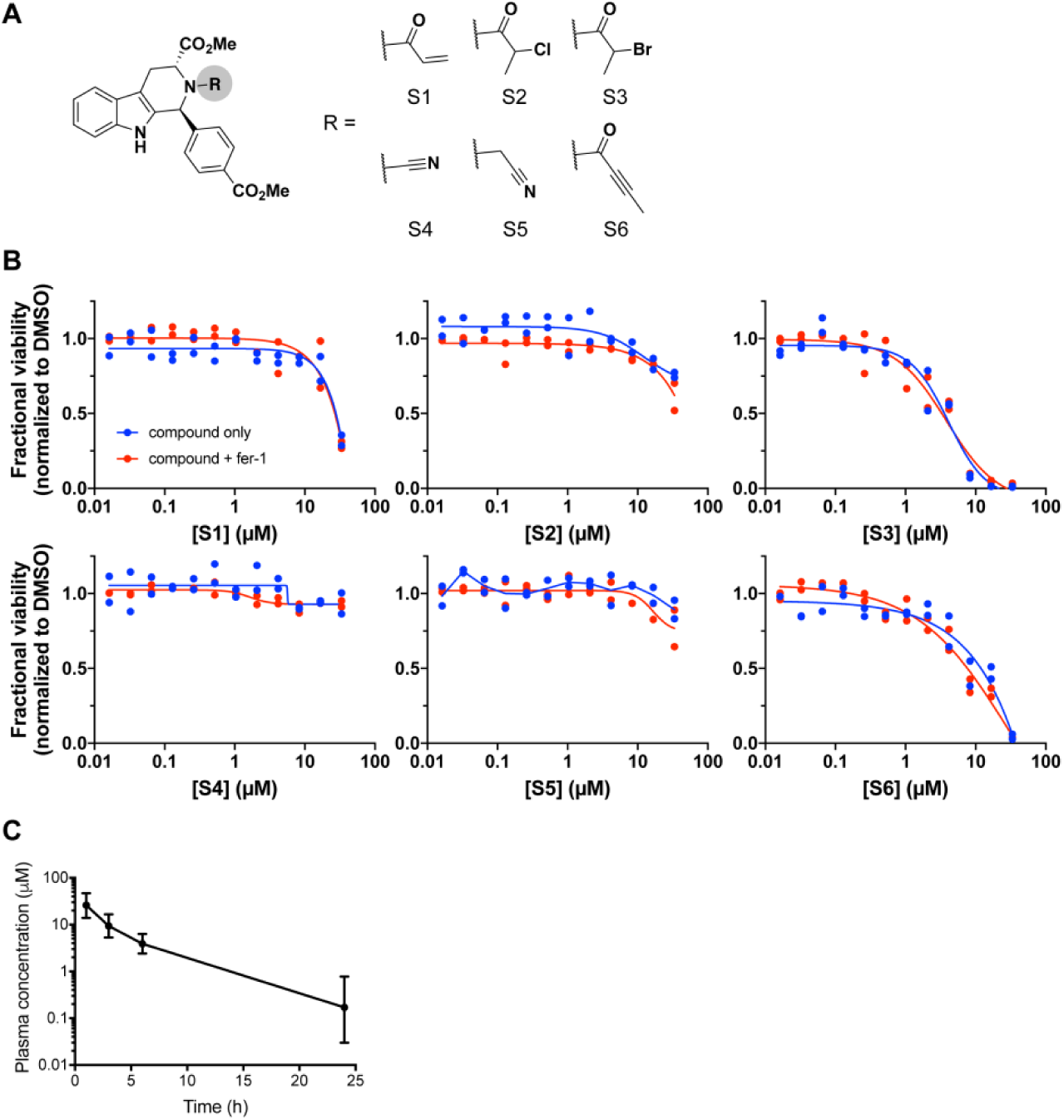
Chloroacetamide GPX4 inhibitors have poor properties that cannot be overcome by substitution of the chloroacetamide group, related to Table 1. (A) Structures of RSL3 analogs with chloroacetamide group substituted with attenuated electrophiles. Data are plotted as individual replicates from two technical replicates. (B) Cellular activity of RSL3 analogs in (A) is not rescued by fer-1 co-treatment. These observations are consistent with a non-overlapping set of reported RSL3 analogs with alternate warheads, including fluoroacetamide, azidoacetamide, and vinylsulfonamide^4^. (C) *In vivo* PK assessment of JKE-1674 in SCID mice. Data are plotted as mean ± s.d., n = 4 biologically independent samples.

**Figure S10.**
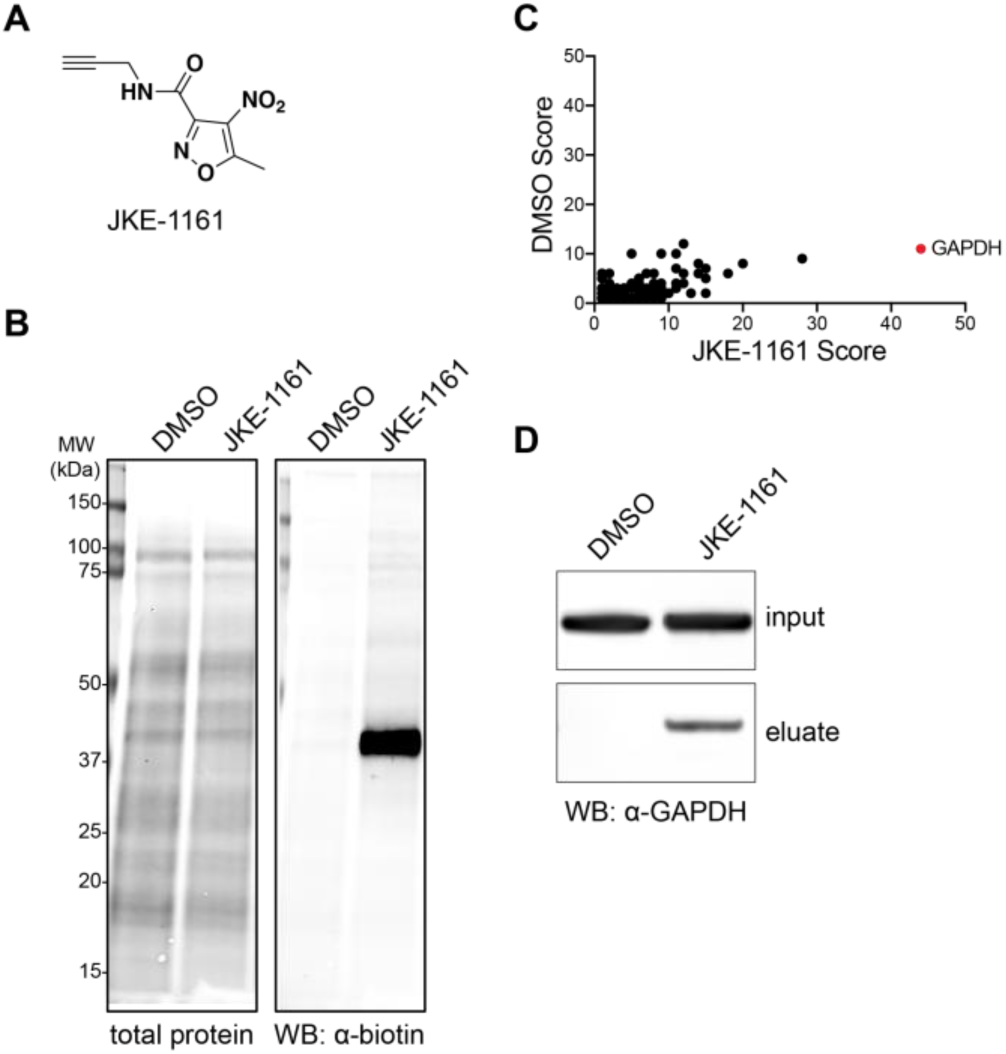
Extensibility of selective covalent interactions by masked nitrile oxides. (A) Structure of a novel nitroisoxazole compound JKE-1161. (B) Assessment of JKE-1161 proteome-wide reactivity reveals strong labeling of a single band at ~40 kDa. (C) Affinity-based chemoproteomics identifies GAPDH as the most enriched target of JKE-1161. (D) Western blot confirmation of GAPDH enrichment by JKE-1161.

## MATERIALS AND METHODS

### Bacterial strains

Origami B (DE3) competent *E. coli* cells were purchased from EMD-Millipore (Billerica, MA) and used for bacterial expression of GPX4. One Shot TOP10 *E. coli* cells were purchased from ThermoFisher Scientific (Waltham, MA) and NEB 5-alpha competent *E. coli* cells were procured from New England BioLabs (Ipswich, MA).

### Mammalian cell lines

HEK293-6E cells used for protein expression were maintained in FreeStyle F17 Expression Medium (ThermoFisher, Waltham, MA). LOX-IMVI (melanoma), CJM (melanoma), WM88 (melanoma), KP4 (pancreatic cancer), and HT1080 (fibrosarcoma) cell lines were procured from the Broad Institute Biological Samples Platform. A498 and CAKI2 renal cell carcinoma cells were procured from ATCC (Manassas, VA). PANC02 (pancreatic cancer) and MC38 (colon cancer) cell lines were a gift from the Dougan Lab (DFCI). All cell lines were maintained according to supplier recommendations except where noted below.

### Cell viability assays

Cell viability experiments were performed by seeding 1000 cells/well in opaque white 384-well plates (Corning). Cells were allowed to adhere for 24 h after which they were exposed to compounds for 72 h. Cellular ATP levels were measured using CellTiter-Glo (Promega) as a surrogate for viability. Ferrostatin-1 (fer-1) rescue experiments were performed using 1.5 µM fer-1 (Sigma Aldrich) added to cells and media at the time of seeding.

### Chemistry and synthesis of chemical materials

Synthesis and analytical characterization of all compounds are detailed in *Synthetic Methods*. Synthetic starting materials, reagents, and solvents were purchased from Sigma-Aldrich, TCI, Acros, Alfa Aesar, AK Scientific Inc., Combi-Blocks Inc., or Enamine Ltd., unless otherwise noted, and were used without further purification.

### Western blot assays

Western blot analyses were performed as described below unless otherwise specified. Lysates were diluted with SDS sample buffer (Boston Bio Products, reducing, 6x) and heated to 95 °C for 5 minutes. Samples (typically 30 μg) were separated by SDS/PAGE (using buffers, gels, and equipment from Invitrogen) and transferred to a nitrocellulose membrane using an iBlot 2 Gel Transfer Device (ThermoFisher). Proteins were visualized with appropriate primary antibodies, IRDye secondary antibodies (LI-COR Biosciences), and an Odyssey imaging system (LI-COR Biosciences).

### Cellular pulldown assay

LOX-IMVI cells were seeded in 6-well plates in RPMI media supplemented with 10% FBS. Alkyne affinity probes were added to the cells and incubated for 1 h at 37 °C (10 mM stocks in DMSO, final alkyne concentration = 10 μM). Media was removed and cells were washed once with ice-cold PBS (pH 7.4). Cells were then lysed in PBS containing 1% Triton X-100 and protease inhibitors (Roche Complete) for 20 min on ice. Samples were cleared by centrifugation (20,000 x g, 4 °C). Total protein content of lysates was assessed with a Bradford assay kit (ThermoFisher, Waltham, MA) and adjusted to 2 mg/mL.

Lysates were subjected to copper-catalyzed azide-alkyne cycloaddition (CuAAC) conditions with azide-PEG3-biotin conjugate (Sigma Aldrich, St. Louis, MO). Typical reactions were performed with a final volume of 120 μL consisting of 100 μL lysate (2 mg/mL; final concentration = 1.67 mg/mL), 2.4 μL SDS (10%; final concentration = 0.2%), 2.4 μL azide-PEG3-biotin conjugate (5 mM in DMSO; final concentration = 100 μM), and 15.2 μL of catalyst mix (final concentration = 1.3 mM Cu_2_SO_4_, 1.3 mM TCEP, and 75 μM TBTA). The catalyst mix stock was prepared by mixing 3 parts TBTA (1 mM in 1:4 DMSO/tBuOH), 1 part 50 mM Cu_2_SO_4_ in water, and 1 part TCEP (50 mM in water, pH 7.0). After addition of all components, CuAAC reactions were vortexed and allowed to react for 1 h at ambient temperature and then diluted with 120 μL 0.2% SDS in PBS. A 40 μL aliquot was removed and quenched with 6x SDS sample buffer as an input control. Pierce high-capacity streptavidin agarose beads (ThermoFisher, Waltham, MA) were added to the remaining sample and rotated overnight at 4 °C. Beads were then separated by centrifugation and washed sequentially with 1% SDS (3 x 1 mL) and PBS (2 x 1 mL). Proteins were eluted by boiling the beads in 75 μL of 2x SDS sample buffer for 10 minutes. The supernatant was removed and analyzed by SDS-PAGE and western blotting.

### Proteome reactivity profiling

Lysates from LOX-IMVI cells treated with alkyne probes were prepared as described above in *Cellular pulldown assay*. CuAAC was used to conjugate samples with IRDye 680RD azide (LI-COR Biosciences, Lincoln, NE). Typical reactions consisted of 100 μL lysate (2 mg/mL; final concentration = 1.67 mg/mL), 2.4 μL SDS (10%; final concentration = 0.2%), 3.6 μL IRDye 680RD azide (1 mM in DMSO; final concentration = 30 μM), and 14 μL of catalyst mix (final concentration = 1.2 mM Cu_2_SO_4_, 1.2 mM TCEP, and 70 μM TBTA). The catalyst mix stock was prepared by combining 3 parts TBTA (1 mM in 1:4 DMSO/tBuOH), 1 part 50 mM Cu_2_SO_4_ in water, and 1 part TCEP (50 mM in water, pH 7.0). After addition of all components, CuAAC reactions were vortexed and allowed to react for 1 h at ambient temperature. The reaction was quenched by addition of 1.2 mL acetone (precooled to −20 °C) and incubated overnight at −20 °C. The sample was then centrifuged (20,000 x g, 10 min, 4 °C) to pellet precipitated proteins. The supernatant was removed and the light blue pellets were washed twice with 600 μL of ice-cold methanol. Pellets were briefly sonicated (using bath sonicator) to fully resuspend contents and were once again centrifuged as described above for 5 minutes following each of the two washes. After the final wash, pellets were allowed to air dry to remove excess methanol. Pellets were suspended in 1x SDS sample buffer, heated at 95 °C for 5 min, and separated by SDS-PAGE. In-gel fluorescence was measured using an Odyssey imaging system (LI-COR Biosciences, Lincoln, NE). Gels were then stained with InstantBlue Coomassie stain (Expedeon, San Diego, CA) and imaged to assess total protein loading.

### Plasmids, expression, and purification of recombinant GPX4 proteins in mammalian cells

All GPX4 expression cassettes used in this manuscript are based on DNA sequence X71973 and encode amino acid sequence M1 to F170 (P36969, with mutations as indicated below). A Kozak sequence precedes the start codon M1, while a stop codon and SECIS element (selencocysteine insertion sequence) appear following codon F170. Three different SECIS elements were tested in the expression cassette: a previously described chimeric element ^57^, the human element from GPX4 X71973-1, and the element from SelN from NM_206926-1. All were found to be equally efficient in expression of GPX4 (U46) when co-expressed with SBP2 (SECIS-binding protein2) in HEK293-6E cells as described below (data not shown).

DNA sequences for the following panel of GPX4 expression cassettes with different mutations and tags were synthesized by GeneArt Technology at Life Technologies. This was achieved using restriction endonuclease sites PstI and BclI at the 5’ and 3’ ends, respectively, and by subcloning into the mammalian expression vector pTT5 ^58^ via the mentioned restriction sites. These DNA constructs and the proteins they encode are designated as detailed below. The final expression vectors, which include additional cassette components, are denoted with the suffix ‘–pTT5.’

GPX4_WT_His_6_, GPX4_U46C_His_6_, GPX4_C66S_His_6_: with C-terminal His6 tag fused in reading frame between F170 and stop codon.

GPX4_WT_Flag, GPX4_U46C_Flag: with an N-terminal Flag tag and a preceding additional start codon fused in-frame to the GPX4 coding region.

DNA encoding rat SBP2 (Q9QX72) contains a Kozak sequence with EcoRI and BamHI sites in the 5’ and 3’ regions, respectively. This was subcloned into expression vector pTT5 to yield the final expression vector, SBP2-pTT5.

Expression plasmids encoding GPX4-pTT5 and SBP2-pTT5 were amplified in *E. coli* (One Shot TOP10) and purified using a QIAprep Minispin Kit (Qiagen; #27104; for small scale) and NuceloBond PC 10000 EF (Macherey-Nagel; #740548) to yield highly pure plasmid preparations suitable for transfection.

For protein expression, a GPX4-pTT5 plasmid together with the SBP2-pTT5 was expressed transiently in HEK293-6E cells. Transfection mixtures ^58^ were prepared by combining a total of 1ug plasmid DNA (comprising both GPX4-pTT5 and SBP2-pTT5 plasmids) for each 1 mL of transfected cell culture with PEI transfection reagent (polyethylenimine, linear, Polysciences # 23966) at a ratio of 1:2 (w/w). This mixture was then added to F17 medium (without any supplements), mixed carefully and incubated for 15 min at room temperature. It was then added to HEK293-6E cells cultured at a density of 1.6 x 10^6^ cells/mL in F17 Medium (Gibco, Invitrogen, Cat# 05-0092DK) supplemented with Pluronic F68 (0.1 %, Gibco # 24040), L-alanyl-glutamine (200mM, Gluta-Max, Invitrogen #25030), G418 (25µg/ml, PAA #P02-012). Cultures were shaken for 72 h at 37 °C in culture vessels ranging in volume from 2 mL to a 10 L bioreactor (Cultibag RM, Sartorius Stedim Biotech). Cells were then harvested by centrifugation (30 min., 1000 x g, 15 °C) and resulting cell pellets were stored at −80 °C.

The purification of recombinant GPX4 proteins expressed in HEK293-6E was performed in two steps using an Äkta Avant chromatography system. This purification procedure involved an initial affinity chromatography step (anti-FLAG column or IMAC, immobilized metal ion affinity chromatography, depending on GPX4 tag) followed by a second size exclusion chromatography step (GE). The pellet from transfected cells was suspended in lysis buffer. A solution of 50 mM sodium phosphate (pH 7.4), 300 mM NaCl, and 0.1% NP-40 was supplemented with protease inhibitor cocktail (Roche Complete) and used for anti-FLAG affinity chromatography. A solution of 50 mM Tris-HCl (pH 7.4), 300 mM NaCl, 10 mM imidazole, 0.1% NP-40, 1 mM DTT, and protease inhibitor tablets (Roche Complete, EDTA-free) was used for IMAC. Suspensions were incubated on ice for 1 h and subsequently centrifuged (63,000 x g, 1 h, 4 °C). The cleared lysate supernatant was used for chromatography.

For IMAC of His-tagged GPX4 constructs, supernatant of the lysate was applied to a Ni-NTA column (Macherey-Nagel; #745400) and washed with buffer 50 mM Tris-HCl (pH 7.4), 300 mM NaCl, 10 mM imidazole, 1 mM DTT. Thereafter, bound protein was eluted from the column using the same buffer supplemented with 300 mM imidazole. For FLAG-tagged constructs, lysate supernatant was applied to a column with anti-FLAG M2 Agarose (Sigma). This was washed with buffer 50 mM sodium phosphate (pH 7.4), 300 mM NaCl and bound protein was eluted using the same buffer supplemented with 250 μg/mL FLAG peptide.

Elution fractions from affinity chromatography were concentrated using Amicon Ultra 15, Centrifugal Filters (10 kDa MW cut-off; Millipore #UFC901024) and applied to size exclusion chromatography columns. The resulting peak fractions were collected, pooled and concentrated again. The buffer used for SEC and final sample preparation was 50 mM sodium phosphate (pH 7.4), 300 mM NaCl, 1 mM DTT for the FLAG-tagged GPX4 protein and 50 mM Tris-HCl (pH 8.0), 150 mM NaCl, 5 mM TCEP for the His-tagged GPX protein. The concentration of final purified products was typically about 1.5 mg/mL and the yield from different GPX4 constructs after final purification was approximately 1 mg product per liter of culture.

### Plasmids, expression, and purification of recombinant GPX4^U^46^C^ protein in bacterial cells

The gene encoding cytoplasmic GPX4 (P36969) was codon-optimized for bacterial *E. coli* expression and fragments were synthesized commercially (GeneArt, Life Technologies). GPX4 fragments were cloned into a derivative of pET21b using the Golden Gate strategy ^59^. The N-terminal 6-histidines tag was followed by two protease cleavage sites (i.e. thrombin and TEV) to facilitate removal of the solubility tag. A C-terminal FLAG-AviTag was used for initial tracking and identification of GPX4 throughout the purification process but later omitted from the construct with a stop codon. Correct insertion of genes into the expression vectors was confirmed by sequencing.

Proteins were expressed in Origami B (DE3) competent cells (EMD-Millipore, Billerica, MA). Large-scale expression of GPX4 proteins was performed by growing cultures in LB at 37 °C with orbital shaking at 180 rpm to mid-log phase (A_600_ ≄ 0.6–0.8). Cultures were then cooled to 18 °C for 30 min before induction with 0.5 mM isopropyl β-D-1-thiogalactopyranoside (IPTG). Induced cultures were grown for 18 h at 18 °C. Cell pellets were harvested via centrifugation and stored at −20 °C.

Cell pellets were lysed in Bugbuster (EMD-Millipore) and Lysonase (EMD-Millipore) by incubating for 15 min at room temperature. Soluble protein was extracted by centrifugation (16,000 x g) and filtered (0.22µM) before loading onto 5 mL Ni-charged HisTrap columns (GE Healthcare). Running buffers for the column were as follows: Buffer A contained 20 sodium phosphate pH 7.4, 500 mM NaCl, 20 mM imidazole, and buffer B contained the same as Buffer A but with 0.5 M imidazole. The column was washed with 3–5 column volumes of buffers A and B, and bound protein was eluted with a linear gradient of imidazole.

Combined fractions were loaded directly onto a preparation grade Superdex 75 10/300 GL column (GE LifeSciences) using an isocratic elution in 25 mM HEPES, 150 mM NaCl, pH 7.4, to remove low-molecular weight contaminants and exchange buffer. Eluted fractions were concentrated and the His(6)-MBP fusion was cleaved by incubation with TEV protease overnight at 4 °C. The next day, digested samples were passed through a Ni^2+^-charged HiTrap column and flow-through were collected. Proteins were concentrated and stored at −80 °C until use.

### Protein crystallography

Wild-type GPX4 (GPX4_WT_His_6_) was crystallized by vapor diffusion using the hanging drop method. Rod-shaped crystals appeared within hours at 20 °C in drops made from 1 µl protein (17.6 mg/ml in 50 mM Tris-HCl (pH 8.0), 150 mM NaCl, 5 mM TCEP) and 1 µl reservoir solution (18-21% PEG 3350 (w/v), 0.1 M MES (pH 6.0), 5% ethanol (v/v)). These reached their final size after 1 day.

The crystal used for the apo structure reported in this manuscript was grown from GPX4 protein that was supplemented additionally with a threefold molar excess of JKE-1674 (from a 100 mM stock solution in DMSO). A 16 h incubation at 4 °C, did not result in modification of any (seleno)cysteine residue. The crystal was briefly immersed in cryo buffer (reservoir supplemented with 15% glycerol) and shock-frozen in liquid nitrogen. Data were collected at 100 K at beamline 14.1 at the Helmholtz-Zentrum Berlin (wavelength λ =0.9184 Å) using a PILATUS detector. Data were processed using the programs XDS ^60^ and XDSAPP ^61^. Data collection statistics are listed in Table S1. The crystal diffracted to a resolution of 1.0 Å and belongs to space group P1 with one GPX4 molecule per asymmetric unit. The structure was solved using Molecular Replacement (program PHASER ^62^ from the CCP4 program suite ^63^) and PDB entry 2OBI as search model and refined using REFMAC5 ^64^. The initial model was rebuilt using the program COOT ^65^, followed by several cycles of refinement and rebuilding. The final refinement statistics are summarized in Table S1.

Initial efforts to generate crystals of GPX4_WT_His_6_ covalently modified with ML162 via incubation (threefold molecular excess, 16 h at 4 °C) followed by crystallization screening failed. An initial screen using 6 commercially available 96-drop crystallization screens and two fine screens around the apo condition described above did not produce any crystals. We suspected that heterogeneous modification of GPX4 by ML162 might have hindered crystallization. We therefore carried out a time-course experiment. Different GPX4:ML162 molar ratios were incubated at three different protein concentrations for 4 h and samples were taken after different time points. The reactions were stopped by addition of trifluoroacetic acid (TFA) and the samples were analyzed by mass spectrometry (data not shown). Based on these results, two reaction conditions were selected for large-scale reproduction. GPX4 (5 mg, 50 µM final concentration) were incubated either for 35 min with a fivefold molar excess of ML162 (condition-1) or for 4h with a 50-fold molar excess of ML162 (condition-2). The reaction mixtures were centrifuged (3 min, 3220 x g) and subjected to size exclusion chromatography (Superdex 75) to separate excess inhibitor and stop the reaction. Peak fractions were concentrated to 16 mg/ml and sitting drops were pipetted using a Mosquito robot (0.2 µl protein solution and 0.2 µl reservoir, 20 °C). MS analysis showed that for condition-1, the protein sample consisted of 50% mono-adduct and 50% double adduct. Condition-2 yielded 100% double-adduct.

Crystals were identified only for the sample from condition-1. These crystals grew in drops made from 20% (w/v) PEG 3350/200 mM magnesium formate and were cryo protected with reservoir supplemented with 15% glycerol. A dataset to 2.3 Å was collected at beamline P11 at PETRA III at DESY, Hamburg (λ =1.0332 Å) using a PILATUS3 6M detector. The crystal belonged to space group *P*2_1_ and diffracted to a resolution of 2.3 Å. The structure was solved using Molecular Replacement ^62^ with the apo structure as search model and refined and rebuilt using REFMAC5 ^64^ and COOT ^65^. The structure contains two GPX4 molecules in the asymmetric unit. Chain B showed extra density on Sec46 and on Cys66 that was interpreted as covalently bound ML162. However, the density was too weak to place the ligand unambiguously.

To address this shortcoming, GPX4_C66S_His_6_ was purified and subjected to a similar MS time-course experiment. Protein (50 µM) was incubated with 125 µM ML162 (pure (*S*)-enantiomer) for 30 min at RT, then centrifuged (3 min, 3220 x g). The reaction was stopped by size exclusion chromatography (Superdex 75). Peak fractions were concentrated to 13.5 mg/ml and sitting drops were pipetted using a Mosquito robot (0.2 µl protein solution and 0.2 µl reservoir, 20 °C). MS analysis showed >95% one-fold modification, <5% free GPX4 and no double adduct. Crystals grew within 1 day with 0.2 M ammonium sulfate and 20% (w/v) PEG 3350 as reservoir solution. They were cryo protected using reservoir solution supplemented with 15 % glycerol. A crystal was mounted at 100 K on a Rigaku 007 diffractometer equipped with a Pilatus 200K detector. Data collection and processing was done using HKL3000 software^66^. The structure was solved using Molecular Replacement ^62^ with the apo structure as search model and rebuilt and refined using COOT and REFMAC5. The crystal belonged to space group *P*2_1_2_1_2_1_ and diffracted to a resolution of 1.54 Å. The structure contains a single GPX4 molecule per asymmetric unit. Clear difference density allowed building of the complete ML162 covalently linked to Sec46. For parameterization, a 3D model of ML162 was generated using Discovery Studio (Dassault Systèmes BIOVIA) and parameter files were generated using software PRODRG ^67^. The final data collection and refinement statistics are summarized in Table S1.

### Preparation of phosphatidylcholine hydroperoxides (PCOOH)

Preparation of PCOOH was accomplished by adapting literature procedures describing lipoxygenase-mediated peroxidation of phosphatidylcholine (PC) ^4,68,69^. Briefly, a solution of 16:0/18:2 PC (100 mg, Avanti Polar Lipids, Inc., Alabaster, AL) in methanol (1 mL) was dissolved in 50 mM borate buffer (170 mL, containing 2.2 w/v% sodium deoxycholate, pH 9.0). Soybean lipoxygenase (2.5 million units; Cayman Chemical, Ann Arbor, MI) was added and the mixture was stirred at room temperature for 30 minutes. The reaction pH was adjusted to 5.0 with 10 N HCl to precipitate deoxycholic acid. The solution was transferred to a separatory funnel and extracted several times with dichloromethane (DCM). The combined DCM fractions were concentrated by rotary evaporation. The crude product was purified by filtration through a short silica plug using a mobile phase consisting of chloroform/methanol/acetic acid/water (100:75:7:4, v/v). Fractions containing PCOOH were identified by TLC with *N,N,N’,N’*-tetramethyl-*p*-phenylenediamine (TPD, Sigma-Aldrich) stain according to a reported protocol ^70^. Combined fractions were dried, dissolved in methanol, and filtered through a 0.2 μm filter. The methanolic PCOOH solution was stored at −20 °C until use.

### GPX4 activity assay

A mass spectrometry-based GPX4 enzymatic activity assay was adapted from a previously described procedure ^4^. LOX-IMVI cells were treated with indicated compounds (10 μM) or DMSO for 1 h at 37 °C. Cells were washed with PBS and lysed by freeze-thaw method (x3) in GPX4 reaction buffer (137 mM NaCl, 2.7 mM KCl, 10 mM Na_2_HPO_4_, 1.8 mM KH_2_PO_4_, 1 mM EDTA, 0.1 mM DFO; pH 7.4). Lysates were cleared by centrifugation (10 min, 20000 x g, 4 °C) and total protein concentration was adjusted to 1.67 mg/mL. Typical enzymatic activity assay mixtures were prepared as follows: 200 μL lysate (1.67 mg/mL in GPX4 reaction buffer), 2 μL of PCOOH in MeOH, and 20 μL GSH solution (100 mM; ≄5 mM final concentration). Reactions were vortexed briefly and incubated at 37 °C for 15 min. Reaction mixtures were then extracted using 250 μL of a 2:1 chloroform/methanol (v/v) solution. Extracts were dried under a stream of nitrogen and reconstituted in methanol before LC-MS/MS analysis.

LC-MS analysis was performed with Acquity RP UPLC system coupled to a XevoG2XS QToF mass spectrometer (Waters). Reconstituted extract was separated on a Waters Acquity RP UPLC BEH-C18 column (2.1 x 50 mm; 1.7 μm particle size) that was maintained at 45 °C. The mobile phase consisted of 10 mM aqueous ammonium acetate (solvent A) and 95:5 acetonitrile/10 mM ammonium acetate (solvent B). The total run time was 8 minutes. UPLC eluate was introduced into the mass spectrometer by positive mode electrospray ionization. Source settings were 120 °C, 50 V cone voltage, 1 kV capiliary voltage, 500 °C desolvation temperature, and 1100 L/h desolvation gas flow. Mass spectrometry experiments were performed in sensitivity mode with a resolution of 20,000 and a mass accuracy of <1 ppm. The lockmass (Leu-Enk, m/z 556.2771) was infused continuously at 5 μL/minute and sampled every 15 seconds. MassLynx (Waters) was used for analysis of mass spectra, compound identification, and chemical formula confirmation analysis. Quantitation was performed using TargetLynx software (Waters).

### Metabolite identification

Time-dependent cellular transformation of ML210 was investigated in LOX-IMVI cells. Cells were cultured in suspension in Williams’ medium E without FCS for up to 24 h, at a cell density of 1×10^6^ cells/mL. Reactions were started by adding 10 µM ML210 to the cell suspension. Aliquots were stopped at pre-determined time points (0, 2, 6, 24, and 48h) by addition of acetonitrile and subsequent centrifugation of precipitated proteins. Supernatants were analyzed by HPLC-DAD-HRMS detection to generate metabolite profiles. Metabolite structures were elucidated by HPLC-MS/MS. Metabolite identification experiments with cells treated with JKE-1674 and JKE-1708 were performed as described for ML210.

### In-cell GPX4 mass spectrometry binding assay

HEK293-6E cells were transfected with either GPX4_WT_Flag-pTT5/SPB2-pTT5 or GPX4_U46C_Flag-pTT5/SPB2-pTT5 in 24-well plates using 2 mL cultures per well. Cells were harvested 72 h following seeding and compounds were added 1, 4 or 24 h before cell harvest. For each time-point, compounds were added at the following concentrations: 1, 5, 10, and 25 µM. Viability of transfected and compound-treated cells was monitored. Cells were harvested by centrifugation, lysed and GPX4 was purified by anti-FLAG chromatography as described above. Purified GPX4 protein was analyzed by denaturing MS and PAGE/Coomassie staining to detect adduct formation.

### Biochemical GPX4 mass spectrometry binding assay

10 µM recombinant N-terminal FLAG-tagged GPX4_WT or GPX4_U46C mutant protein was incubated with 100 µM compound (1% v/v final DMSO concentration) at room temperature for 2 h. The reaction was quenched by adding 1 µL 5% v/v TFA to 15 µL reaction volume and was subjected to LC-MS analysis. LC-MS analysis of covalent binding was performed with a Waters SYNAPT G2-S quadrupole time-of-flight mass spectrometer connected to a Waters nanoAcquity UPLC system (Waters, Milford, MA). Samples were loaded on a 2.1 x 5 mm mass prep C4 guard column and desalted with a short gradient (3 min.) of increasing concentrations of acetonitrile at a flow rate of 100 µL/min. Spectra were analyzed by using MassLynx v4.1 software (Waters, Milford, MA) and deconvoluted with the MaxEnt1 algorithm.

### Lipid peroxidation assay by flow cytometry

U2OS cells were seeded at 15,000 cells per well in 96-well plates. After 48 h, culture media was replaced with 200 μL media containing either +/-ferrostatin-1 and DMSO or the indicated GPX4 inhibitor (10 µM). Cultures were incubated at 37°C for 2 h. 30 minutes before the end of the incubation period, 10 µM BODIPY 581/591 C11 (MOLECULAR Probes #C10445) was added to cells. Cells were harvested in 200 μL PBS + 0.1% BSA and subjected to flow cytometry analysis to examine oxidation of BODIPY 581/591 C11, a reporter for lipid ROS levels in cells. A BD FACSCanto II instrument was used for the flow cytometer analysis.

### Lipid peroxidation assay by imaging

LOX-IMVI cells were seeded at 5,000 cells per well in a CellCarrier Ultra 96-well plate (Perkin-Elmer, Waltham, MA) in 150 μL of RPMI medium supplemented with 10% FBS, 1% pen-strep, 5 μg/mL plasmocin, and 1 µM fer-1 (where indicated). Cells were allowed to adhere for 24 h at 37 °C and then treated with the indicated compounds or DMSO for 90 min at 37 °C. During the last 30 min of incubation, 60 nM DRAQ7 (Abcam, Cambridge, MA), 1 μg/mL Hoechst 33342 (ThermoFisher, Waltham, MA), and 1 μM BODIPY 581/591 C11 (ThermoFisher, Waltham, MA) dyes were added to the wells. Cells were imaged using an Opera Phenix High-Content Screening System (Perkin-Elmer, Waltham, MA) equipped with 405 nm, 488 nm, 560 nm, and 647 nm lasers. Image analysis was performed with Harmony High-Content Imaging and Analysis software (Perkin-Elmer, Waltham, MA).

### Generation of cell lines expressing 3xFLAG-GPX4 mutants

Replacement of the GFP sequence in the pBabe-puro GFP-GPX4-cyto expression vector (gift from Stockwell lab) ^4^ with a 3xFLAG sequence (MDYKDHDGDYKDHDIDYKDDDDK) was performed with Q5 Site-Directed Mutagenesis Kit (New England Biolabs, Ipswich, MA). Cysteine (U46C) and alanine (U46A) active-site mutants were obtained by site-directed mutagenesis of codon TGA to TGC and GCC, respectively.

### Compound solubility determination

The high-throughput screening method to determine aqueous drug solubility was performed as described previously by Onofrey, et al (Lit. No. AN1731EN00; 2003)

Test compounds were applied as 1 mM solutions in DMSO. After the addition of PBS buffer (pH 6.5), the solution was shaken for 24 h at RT. Undissolved material was separated by filtration. The compound dissolved in the supernatant was diluted accordingly and quantified by HPLC-MS/MS.

### Compound thiol reactivity assay

Stability of compounds to thiols was determined by HPLC-UV according to a reported procedure ^71^. Reactions were performed in the presence of a 100-fold excess of thiol.

Compound stock solutions in DMSO (5 µL, 10 mM) were dissolved in 1 mL acetonitrile. Cysteine and glutathione, respectively, were dissolved in PBS buffer (pH 7.4)/acetonitrile 7:3 to give 500 µM solutions. 100 µL of drug solution were added to 1 mL thiol reaction solution. Reactions were analyzed by HPLC-Uvc immediately after mixing for time zero injection and then again after 1, 2, 4, and 24 h. Incubations were performed at 37 °C. Degradation rates (recovery in %) were calculated by relating peak areas after 1, 2, 4, and 24 h to the time zero injection peak area.

### Compound pH stability assay

Solution stability of compounds was determined by HPLC-UV. Compound stock solutions in DMSO (5 μL, 10 mM) were diluted in acetonitrile (1 mL), and 100 μL of the resulting solution was transferred to 1 mL of the respective reaction buffer (pH 10, 7, and 1). After mixing thoroughly, the reactions were incubated at 37 °C. Samples were monitored by HPLC-UV immediately after mixing (t=0) and then again after 1, 2, and 24 h. The percent recovery of a compound was calculated by relating the HPLC-UV peak at 1, 2, and 24 h to the time zero injection.

### Compound plasma stability assay

The stability of compounds in human (P9523 Sigma), rat (Charles River, Wilmington, MA), and mouse plasma was determined by HPLC-MS/MS. N-(4-Chlorophenyl)-2-[(pyridin-4-ylmethyl)amino]benzamide (CAS 269390-69-4, SIGMA) was used as an internal standard.

Compound stocks solutions were prepared in DMSO (30 μM), 5 μL of which was added to 495 μL of plasma. Compound-plasma samples were incubated at 37 °C and then homogenized using an Eppendorf Mixmate (1200 rpm, 5 min). The reaction was quenched by transferring 50 μL of the sample to 200 μL of internal standard solution, followed by homogenization (Eppendorf Mixmate, 1200 rpm, 5 min) and centrifugation (4000 rpm, 15 min, 4°C). For initial time point (t = 0 min) samples, the reaction was quenched immediately after the compound stock solution was added to plasma. Further measurements were taken after 1, 2, and 4 h. Peak areas of 1, 2, and 4 h measurements were compared to initial (t = 0 min) peak to determine percent recovery.

### Determination of compound logD values

Reversed-phase high performance liquid chromatography (HPLC) was used to determine compound distribution coefficient (logD) values at pH 7.5 using a reported method for determining hydrophobicity constants ^72^. Test compound stock solutions (0.67 mM in DMSO) were dissolved in acetonitrile/water (1:1, v/v) and analyzed by HPLC. A homologous series of n-alkane-2-ones (C3-C16, 0.02 M in acetonitrile) was used for calibration. Compound lipophilicity was determined by comparison to the calibration curve.

### Metabolic stability assessment in rat hepatocytes

Rat liver cells cultured in Williams’ medium E containing 5% fetal calf serum (FCS) were aliquoted into glass vials at a density of 1.0 × 10^6^ vital cells/mL. Test compounds were added to a final concentration of 1 μM. During incubation, hepatocyte suspensions were shaken continuously. Aliquots were taken at 2, 8, 16, 30, 45, and 90 min, to which an equal volume of cold MeOH was added immediately. Samples were frozen at −20 °C overnight then centrifuged (15 min, 3000 rpm). The supernatant was analyzed using an HPLC system with LC-MS/MS detection.

The half-life of a test compound was determined from the concentration–time plot. From half-life, the intrinsic clearance was calculated. The well-stirred liver model was used to calculate hepatic *in vivo* blood clearance (CL) and maximal oral bioavailability (Fmax). The following parameters were used for calculations: liver blood flow, 4.2 L/h/kg; specific liver weight, 32 g/kg body weight; liver cells *in vivo*, 1.1 × 10^8^ cells/g liver; liver cells *in vitro*, 1 × 10^6^/mL.

### Metabolic stability assessment in liver microsomes

The *in vitro* metabolic stability of test compounds was determined by incubating them at 1 μM with a suspension of liver microsomes in 100 mM PBS (pH7.4) and at a protein concentration of 0.5 mg/mL at 37 °C. Microsomes were activated by adding a cofactor mix containing 8 mM glucose-6-phosphate, 4 mM magnesium chloride; 0.5 mM NADP and 1 IU/ml glucose-6-phosphate-Dehydrogenase in PBS (pH 7.4). The metabolic assay was started shortly afterwards by adding the test compound to the incubation at a final volume of 1 mL. Organic solvent in the incubations was limited to ≤0.01% dimethylsulfoxide (DMSO) and ≤1% acetonitrile. During incubation, the microsomal suspensions were shaken continuously and aliquots were taken at 2, 8, 16, 30, 45 and 60 min, to which equal volumes of cold methanol were immediately added. Samples were frozen at −20 °C overnight, subsequently centrifuged for 15 minutes at 3000 rpm and the supernatant was analyzed using an HPLC-system with LCMS/MS detection.

The half-life of a test compound was determined from the concentration-time plot. From half-life, the intrinsic clearance was calculated. The well-stirred liver model was used to calculate hepatic *in vivo* blood clearance (CL) and maximal oral bioavailability (Fmax). The following parameters were used for calculations: liver blood flow – 1.32 L/h/kg (human), 5.4 L/h/kg (mouse); specific liver weight – 21 g/kg (human), 43 g/kg (mouse); microsomal protein content – 40 mg/g liver.

### Caco-2 permeability assay

Caco2 Drug Permeability Assay. Caco-2 cells (purchased from DSMZ Braunschweig, Germany) were seeded at a density of 4.5 × 10^4^ cell per well on 24-well insert plates, 0.4 μm pore size, and grown for 15 days in DMEM medium supplemented with 10% fetal bovine serum, 1% GlutaMAX (100x, GIBCO), 100 μg/mL penicillin, 100 μg/mL streptomycin (GIBCO), and 1% nonessential amino acids (100x). Cells were maintained at 37 °C in a humified 5% CO2 atmosphere. Medium was changed every second or third day. Before running the permeation assay, the culture medium was replaced with an FCS-free HEPES-carbonate transport buffer (pH 7.2). For assessment of monolayer integrity, the transepithelial electrical resistance (TEER) was measured. Test compounds were dissolved in DMSO and added either to the apical or basolateral compartment in a final concentration of 2 μM. Before and after incubation at 37 °C, samples were taken from both compartments. Analysis of compound content was performed following precipitation with methanol and LC-MS/MS analysis. Permeability (P_app_) was calculated in the apical to basolateral (A→B) and basolateral to apical (B?A) directions. The apparent permeability was calculated using following equation:

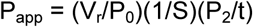

where V_r_ = volume of medium in the receiver chamber, P_0_ = measured peak area of the test drug in the donor chamber at t = 0, S = surface area of the monolayer, P_2_ = measured peak area of the test drug in the acceptor chamber after incubation for 2 h, and t = incubation time. The efflux ratio basolateral (B) to apical (A) was calculated by dividing P_app_ (B-A) by P_app_ (A-B). In addition, the compound recovery was calculated. Reference compounds were analyzed in parallel as assay controls. All samples were analyzed by LC-MS/MS.

### Cellular thermal shift assay (CETSA)

For intact-cell CETSA experiments, cells were pretreated with 10 μM compound or DMSO control (0.1%, v/v) for 1 h at 37 °C. Media was then aspirated and cells were washed with PBS (pH 7.4). Adherent cells were detached from the flask with trypsin-EDTA and pelleted by centrifugation (5 minutes at 500 x g). Cells were aliquoted into PCR tubes (50 μL volume, ≄1 million cells/condition) for heating at different temperatures (typically 40-67 °C in 3 °C increments) in a thermal cycler for 3 minutes. Samples were allowed to cool to room temperature for an additional 3 minutes. Cells were lysed by either three freeze-thaw cycles in liquid nitrogen, or by the addition of Triton X-100 solution (1% final TX-100 volume, PBS pH 7.4) and subsequent incubation on ice for 20 minutes with occasional vortexing. After lysis, cells were centrifuged (20 minutes at 20,000 rcf, 4 °C) to remove insoluble material. The soluble fraction was carefully separated and diluted with 6x SDS loading buffer for SDS-PAGE and western blotting analysis. Three independent replicates were performed typically.

For lysate CETSA experiments, lysate from untreated cells was prepared as described above and diluted in PBS (pH 7.4) to a total protein concentration of 1 mg/mL. Samples were treated with 10 μM compound (or 0.1% v/v DMSO control) for 1.5 h at 37 °C. After compound treatment, samples were aliquoted (50 μL per sample) into PCR tubes. The remaining sample preparation steps were performed as described above for CETSA with intact cells.

### Animal studies

All animal experiments were conducted in accordance with the European animal welfare law and approved by local authorities.

#### Pharmacokinetic (PK) studies in mice

To determine the pharmacokinetic properties of JKE-1674 in SCID mice (Janvier Labs) after oral administration, blood samples for analysis of plasma were taken from three animals/time point at 1, 3, 6 and 24 h after single oral dose of JKE-1674 at 50 mg/kg formulated in PEG400/Ethanol (90/10, v/v). Blood was collected into tubes containing lithium heparin, centrifuged to isolate plasma, precipitated with acetonitrile (1/5, v/v) and analyzed by LC-MS/MS.

#### Toxicity assessment in mice

Tolerability of JKE-1674 was assessed preceding pharmacokinetic measurements. Over a period of 7 consecutive dosing days, average body weight loss in mice which received 50 mg/kg active compound did not exceed 10%. Doses higher than 50 mg/kg were not tolerated in this experimental setting.

